# IS200/IS605 Family-Associated TnpB Increases Transposon Activity and Retention

**DOI:** 10.1101/2022.10.12.511977

**Authors:** Davneet Kaur, Thomas E. Kuhlman

## Abstract

The IS200/IS605 family of insertion sequences are abundant mobile elements associated with one of the most numerous genes found in nature, *tnpB*^1–3^. Previous studies suggest that TnpB protein may be an evolutionary precursor to CRISPR Cas enzymes, and TnpB has received renewed interest having itself been shown to function as a Cas-like RNA-guided DNA endonuclease^3,4^. However, interpretation of the fundamental role of TnpB in transposition and how it contributes to genome dynamics^5^ remains controversial without direct, real-time measurement in live cells. Here, using a suite of fluorescent reporters coupled to transposition in live *Escherichia coli*, we show that IS608-TnpB causes increased transposon activity, and assists in preventing transposon loss from host genomes. Analyzing our results through a mathematical model of transposon dynamics, we discuss the multifaceted roles it may play in transposon regulation. The mutually beneficial transposon-TnpB interaction may explain the prevalence of *tnpB*, creating conditions for the appropriation of TnpB’s RNA-guided endonuclease activity for adaptive immunity.

**SIGNIFICANCE STATEMENT:** Phylogenetic evidence suggests that *tnpB*, one of the most numerous genes found in nature, is the ancestral form of CRISPR-Cas enzymes and played a critical role in the evolution of adaptive immunity. However, the role TnpB plays in transposition that has contributed to its wide distribution remains unclear. Here, we use a unique approach that couples fluorescent reporters to transposition to non-perturbatively quantify transpositional dynamics in live cells. In contrast to previous indirect methods suggesting that TnpB suppresses transposition, our results instead clearly demonstrate that TnpB significantly increases transposition rates and enhances transposon retention within the host genome, resulting in a mutually beneficial interaction between transposons and TnpB that can account for its wide distribution.

## INTRODUCTION

Transposable elements (TEs) are mobile DNA sequences that can move around or create copies of themselves in a host organism’s genome^6^. TEs have been established as major contributors to disease, development, and evolution^7–16^. As such, revealing their network of interactions and dynamics is a key factor in understanding the dynamics of evolution. Together with DNA modification systems and auxiliary genes, TEs form a tangled web of interactions that are integral to the proper functioning of organisms^14,17–19^. In some cases, TE activity is even demonstrably necessary for the viability of an organism^20^. Yet, our understanding of the scope of TE interactions and dynamics remains limited.

Despite their ubiquity and importance, limitations of existing experimental technologies have hindered our understanding of TE activity and interactions in living cells. Many existing studies attempt to extrapolate transposition rates from phylogenetic comparisons of related species^21,22^, or from endpoint analyses of the expression of some reporter gene coupled to transposition^5^. However, extrapolation of dynamic rates from such static methods would require exhaustive knowledge of mechanistic models of transposition, making them inappropriate for exploratory studies into the effects of incompletely characterized systems. To begin to remedy these difficulties, we have developed a suite of fluorescent reporters coupled to transposition to study the bacterial TE IS608^23^. These fluorescent reporters allow us to measure transpositional dynamics within the simple environment of *Escherichia coli*, an ideal host to study the interaction of IS608 with uncharacterized auxiliary proteins.

IS608, originally from *Helicobacter pylori*, is representative the of IS200/IS605 family of TEs. The IS200/IS605 family of TEs all transpose through similar ‘peel-and-paste’ mechanisms and are widely distributed, with 153 members spread over 45 genera and 61 species of eubacteria and archaea^1^. IS608 is an autonomous TE, containing all genetic features required for transposition, and in particular codes for the transposase TnpA that executes recognition, excision, and reintegration^25–30^. In addition to *tnpA*, IS608 contains an additional gene of much recent interest but thus far unclear function, *tnpB*^5,31,32^.

TnpB proteins are an extremely abundant family of nucleases that are encoded by many bacterial and archaeal TEs^1,2^ which often contain only *tnpB*^33^. More than 1 million putative *tnpB* loci have been recently identified in bacterial and archaeal genomes, making it one of the most common prokaryotic genes^3^. Furthermore, sequence similarities and structural homologies suggest that TnpB may be a possible ancestor of Cas12 and IscB^34^, which is a putative ancestor of Cas9^1,34,35^. Recently, TnpB has been shown to function as a programmable RNA-guided endonuclease, like the Cas proteins, that under different conditions will make either dsDNA or ssDNA breaks^3,4^. TnpB’s abundance and conserved association with transposons are indications of its importance in the regulation of TEs and in the evolution of prokaryotic systems, and its programmability and diversity of function suggests that TnpB may be a promising candidate for development as a genome editing tool.

However, the role of TnpB in transposition remains poorly understood. Previous studies attempting to determine the role of TnpB in transposition have, for example, employed measurements coupling IS200/IS605 transposition to the development of antibiotic resistance measured at a single time point after long periods of growth^5^. By measuring the number of bacteria with antibiotic resistance at a single time point, they have attempted to extrapolate these end-point measurements to suggest that TnpB suppresses the dynamic rates of transposition^5^. However, without exhaustive knowledge of transposition mechanisms, such as the recently proposed homing mechanism where TnpB aids in reinserting a copy of the TE at locations from where it has previously excised^4^, such estimations of dynamic rates using static endpoint measurements are fraught with difficulty. The proposed homing mechanism would, in fact, lead to re-disruption of antibiotic resistance genes reconstituted as a result of standard transposition in the above-described assays, and would consequently result in apparent artefactual suppression of transposition regardless of other alterations to the underlying dynamics. Hence the functional role of TnpB remains controversial.

The limitations exemplified above demonstrate that there is a need for dynamic studies of TE behavior in living cells and the effects of TE activity on those cells. Here, we employ our developed IS608 systems to quantitatively measure the effects of TnpB expression on the dynamics of transposition of IS608 in live cells. We find, contrary to previous static measurements, that TnpB in fact dramatically increases the rates of transposition. Through analysis of these data through a mathematical model of IS608 dynamics, we find that, through enhanced rates of transposition and the previously proposed homing mechanism, TnpB aids in facilitating the retention of TEs within the host genome over time. This mutually beneficial interaction between TEs and IS608 may have set the stage for the vast proliferation of tnpB genes in nature and their eventual adaptation as systems of bacterial adaptive immunity.

## RESULTS

### Measuring TnpA, TnpB, and excision levels

We created an inducible IS608 (Tn4rev, previously referred to as ISLAG^23^) that is tagged with fluorescence reporter genes that allow visualization of TE protein levels and activity (**Fig. 1A.i**. and **Supplementary Methods and Statistical Analysis SIII.i**) and introduced it to *E. coli*. By removing the hairpin structures in Tn4rev, RE and LE, that are recognized by TnpA for excision we also created an immobile version of the TE (**Supplementary Fig. S1A.iv**). In all strains, the *P*_*Ltet-01*_ promoter is induced by addition of anhydrotetracycline (aTc)^36^ to generate TnpA translationally fused to Venus fluorescent protein. The use of this inducible promoter allows for simple and precise control of TnpA levels within individual cells^23^. On the plasmid, the TE splits the −10 and −35 sequences of a strong constitutive *P*_*LacIQ1*_ promoter^37^ for the expression of the blue reporter *mCerulean3*^38^. This promoter generates Cerulean fluorescent protein upon successful TE excision. We have found in previous work that excision results in a short burst of Cerulean fluorescence followed by decay^23,39^. Hence, Cerulean fluorescence reports the rate of excision, and Cerulean fluorescence integrated over time (“Cumulative Cerulean”) corresponds to the number of excision events; see **Supplementary Methods and Statistical Analysis SIII.ii** and **Supplementary Fig. S7** for details.

**Figure 1:**
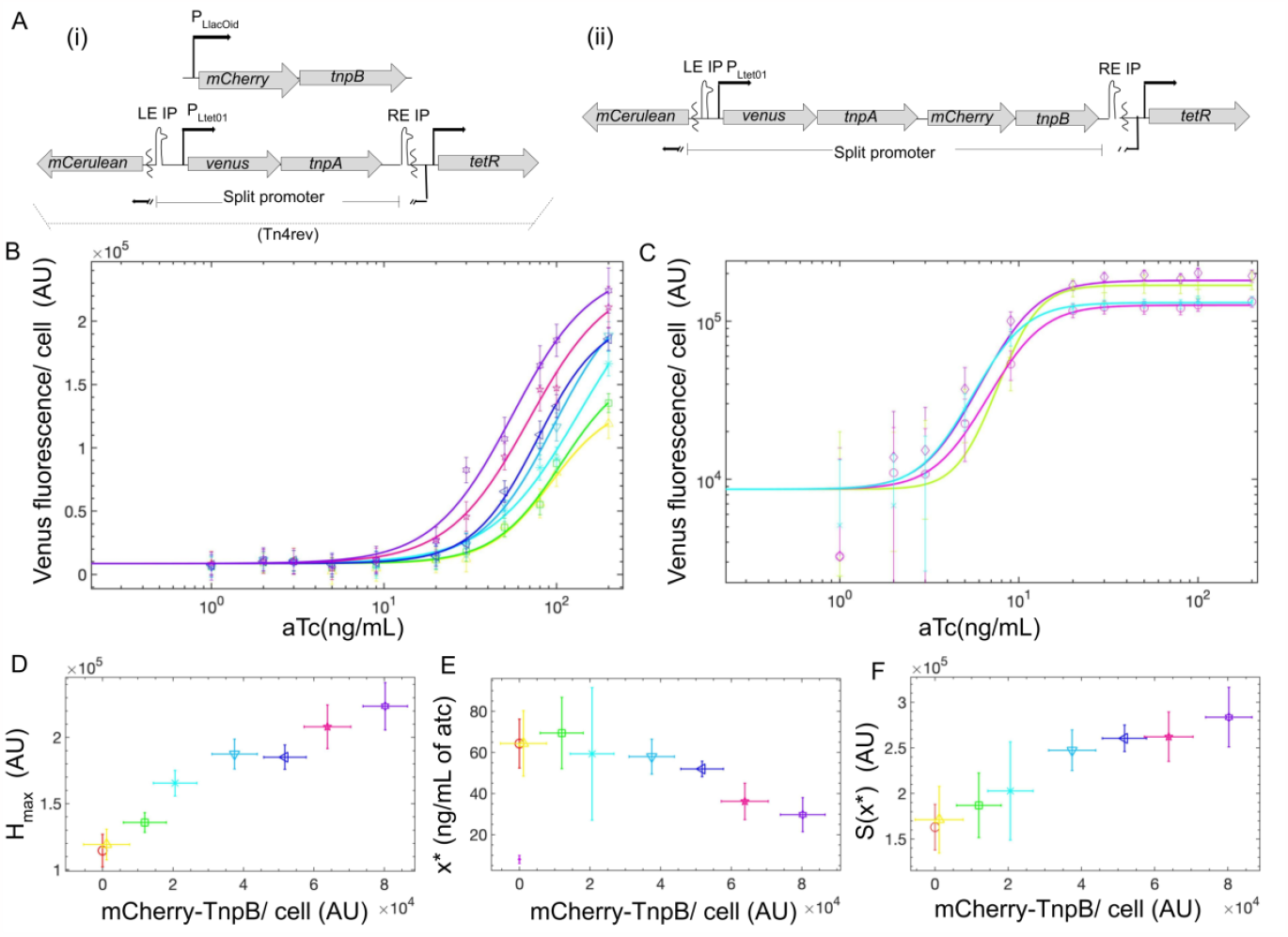
DESIGN. **(A) Genetic Constructs for Inducible Transposon Quantification:** Transposons constructed for experiments. (i) TnpB introduced *in trans* with Tn4rev, MG1655 ∆*lac* pJK14-*P*_*Ltet-01*_*-Tn4rev &* pZA31-P_*LlacOid*_-*mCherry-tnpB* and (ii) *in cis*, MG1655 ∆*lac* pJK14-P_*Ltet-01*_-*tn4rev-mCherry-tnpB &* pZA31-P_*LlacOid*_-SmR. **(B)** Venus fluorescence data as a function of aTc concentration for when TnpB is introduced *in trans* with Tn4rev for different [IPTG] [0μM (yellow, ∆), 10μM (green, □), 20μM (turquoise, ✳), 50μM (teal, ▽), 100μM (navy, ◁), 200μM (magenta, 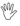) & 2000μM (violet, ✵)]. **(C)** (i) Immobilized transposon, no IPTG (purple, ◊), (ii) Immobilized transposon, with 2000μM IPTG (lizard green, +), (iii) Immobilized transposon with TnpB, no IPTG (fuchsia, ◯) and (iv) Immobilized transposon with TnpB, with 2000μM IPTG (arctic, X). **(B-C)** Points are fit to a Hill function (solid lines, same color; see **Supplementary Information SI.i**). Quantitative features of the response are extracted and included in **Supplementary Table S1** and plotted in **D-F** with corresponding color and symbols. Also included is the Tn4rev without TnpB strain: MG1655 ∆*lac* pJK14-*P*_*Ltet-01*_-Tn4rev & pZA31-P_*LlacOid*_-SmR (red, **◯**) **(D)** TnpB induction results in higher saturating TnpA-Venus (*H*_*max*_). Peak Venus fluorescence is a relative measurement of the population-averaged number of transposons per cell. **(E)** Increased TnpB induction results in earlier TnpA-Venus expression [inflection point of curves in (**B**), *x\**]. **(F)** TnpB induction results in increased TnpA-Venus sensitivity to aTc [slope at inflection point *S(x*)*].

As many other TE-associated proteins exhibit strong *cis*-preference^40^, we introduced *tnpB* to this system in both *trans* and *cis* (**Figs. 1A.i&ii** respectively). For the *trans* combination, *P*_*LlacOid*_*-mCherry-tnpB* is independently induced with isopropyl β-d-1-thiogalactopyranoside (IPTG) (**Supplementary Information SI.i)**. In **Fig. 1A** both strains have *mCherry* (red) translationally fused to *tnpB*, providing a measure of TnpB levels. Control strain lacking mCherry fusion to *tnpB* (**Supplementary Fig. S1A.v**) confirms that this fusion does not affect the activity of TnpB. Furthermore, for strains lacking TnpB, we express the protein AadA from the same pZA31 plasmid and *P*_*LlacOid*_ promoter used for TnpB expression to control for effects of general protein expression on growth and physiology (pZA31-P_*LlacOid*_-SmR). That the Venus-TnpA fusion does not significantly affect activity of TnpA was confirmed in previous work.^23^

### TnpB tunes TnpA concentration to increase excision

Venus-TnpA concentration, excision rate, and the number of plasmid-excision events increase with TnpB (**Figs. 1B,1D,2A,2C**, and **Supplementary Fig. S2,S3A-C,S4**) and become more sensitive to induction with aTc (**Figs. 1E-F,2D-E, Supplementary Fig. S3D-E**, and **Supplementary Information SI.ii-iv**). Excision of the TE happens in response to the TnpA concentration in cells. Hence, we quantify the excision response (cumulative Cerulean fluorescence) as a function of Venus-TnpA concentration for all concentrations of TnpB. As the concentration of transposase increases, the number of excision events increases proportionally (**Fig. 2B**). The data collapse to a single curve, suggesting that TnpB does not directly affect excision. Rather, TnpB expression causes higher concentrations of TnpA, which then catalyzes greater excision (**Fig. 2B, Supplementary Fig. S5**, and **Supplementary Information SI.v**). A linear regression is fit to the cumulative Cerulean data upon initiation of Venus-TnpA expression to find a slope of 545.79 AU with an R^2^ value of 0.913 (**Fig. 2B**).

**Figure 2:**
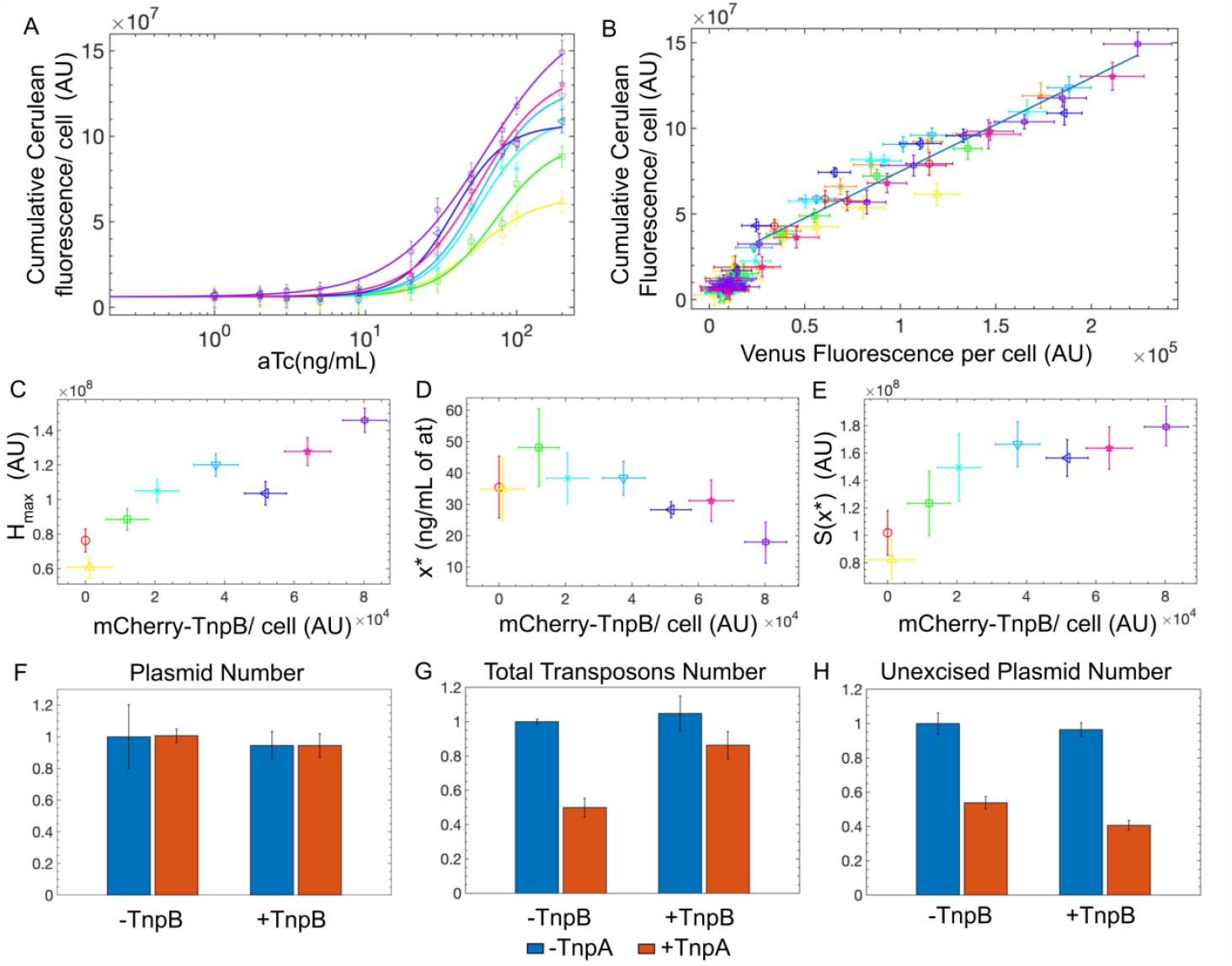
EXCISION AS A FUNCTION OF TNPA. **Transposon Activity and Retention in Response to TnpB. (A)** Cumulative Cerulean fluorescence data as a function of aTc concentration (0ng/mL, 1ng/mL, 2ng/mL, 3ng/mL, 5ng/mL, 9ng/mL, 20ng/mL, 30ng/mL, 50ng/mL, 80ng/mL, 100ng/mL and 200ng/mL) for TnpB *in trans* with Tn4rev for different IPTG concentrations [0μM (yellow, ∆), 10μM (green, □), 20μM (turquoise, ✳), 50μM (teal, ▽), 100μM (navy, ◁), 200μM (magenta, 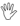) & 2000μM (violet, ✵)]. Cumulative Cerulean fluorescences are fit to a Hill function (solid lines, same color; see **Supplementary Information SI.i**). Quantitative features of the responses are extracted and included in **C-E** and **Supplementary Table S2. (B)** Number of excision events is proportional to transposase concentration: Cumulative Cerulean fluorescence vs. Venus-transposase concentration is plotted for Tn4rev only strain (red, **◯**), TnpB introduced *in trans* with Tn4rev for all concentrations of IPTG (same colors and symbols as in **A**), and TnpB introduced *in cis* with Tn4rev (orange, X). The data collapse to a single curve and a linear regression is fit to the data (solid line). **(C)** TnpB induction results in higher saturating Cumulative Cerulean levels (*H*_*max*_). **(D)** Increased TnpB induction results in earlier Cumulative Cerulean-excision [inflection point of curves in **A**, *x\**]. **(E)** TnpB induction results in increase excision sensitivity [slope at inflection point of curves in **A** *S(x*)*] to aTc concentration. **(F-H)** qPCR reactions determining relative numbers for (**F**) plasmid, (**G**) total transposon, and (**H**) unexcised plasmid count. P-values quantifying the significance of differences are given in **Supplementary Tables S4 and S5**. See **Supplementary Table S7** for primer sequences.

Introduction of TnpB to the immobilized transposon does not result in an increase in Venus-TnpA (**Fig. 1C**), suggesting that TnpB is affecting transposition and not TnpA stability to achieve higher Venus-TnpA concentrations for the active transposon strains. This suggests that TnpB is increasing the number of transposons per cell, either through better retention of transposons or replication.

### TnpB increases transposon excision and retention

*E. coli* with TnpB introduced *in trans* with Tn4rev were grown with and without TnpA induction (0 or 200ng/mL aTc) and with and without TnpB induction (0 or 2000μM IPTG). qPCR reactions (described in **Methods**) were performed to measure relative quantities per cell of plasmid numbers, total transposon numbers, and plasmid-borne transposon numbers (**Supplementary Methods and Statistical Analysis SIII.iii-iv**). We find that the total plasmid copy number per cell is consistent for all concentrations of inducers (**Fig. 2F**). The total transposon number drops when cells are grown with TnpA induction alone relative to when transposition is not induced (**Fig. 2G**, left set of bars). Simultaneous induction with TnpB improves transposon retention at values closer to when no transposition is induced (**Fig. 2G**, right set of bars). Induction of TnpB alone has no significant effect on transposon count (**Fig. 2G, Supplementary Table S4**). Finally, the number of transposons remaining in the original plasmid number drops when cells are grown with TnpA induction alone relative to when transposition is not induced (**Figure 2H**, left set of bars). Simultaneous induction with TnpB further reduced plasmid-transposon numbers, indicating an increase in excision with TnpB (**Fig. 2H**, right set of bars). Induction of TnpB alone has no significant effect on plasmid-transposon count (**Fig. 2H, Supplementary Table S5**).

### TnpB increases the effects on growth from each transposition event

As previously noted^4^, TnpB is unstable in the absence of other transposon features, which we note here for the strain with TnpB alone: MG1655 ∆*lac* pJK14 pZA31-P_*LlacOid*_-*mCherry-tnpB* (**Fig. 3A**, red), which achieves significantly lower mCherry-TnpB concentrations compared to the strain with TnpB introduced *in trans* with Tn4rev for the same range of IPTG induction (**Fig. 3A**, blue). Consequently, lower concentrations of TnpB do not cause significant growth defects. When TnpB is expressed with an active TE, we observe higher concentrations of TnpB and growth defects (**Fig. 3A**, blue), possibly due to TnpB’s dsDNA endonuclease activity. Upon the co-expression of TnpA and TnpB for the immobile TE strain, there is greater growth defect than for the immobile TE without TnpB expression for the same concentrations of TnpA (**Fig. 3B**), suggesting that TnpA and TnpB have an undetermined collaborative interaction that is independent of the execution of transposition. We will determine if TnpA and TnpB colocalize and interact within their hosts in future work as this could be a promising direction towards developing a TnpA-TnpB-based genomic editing tool capable of making large genomic insertions.

**Figure 3:**
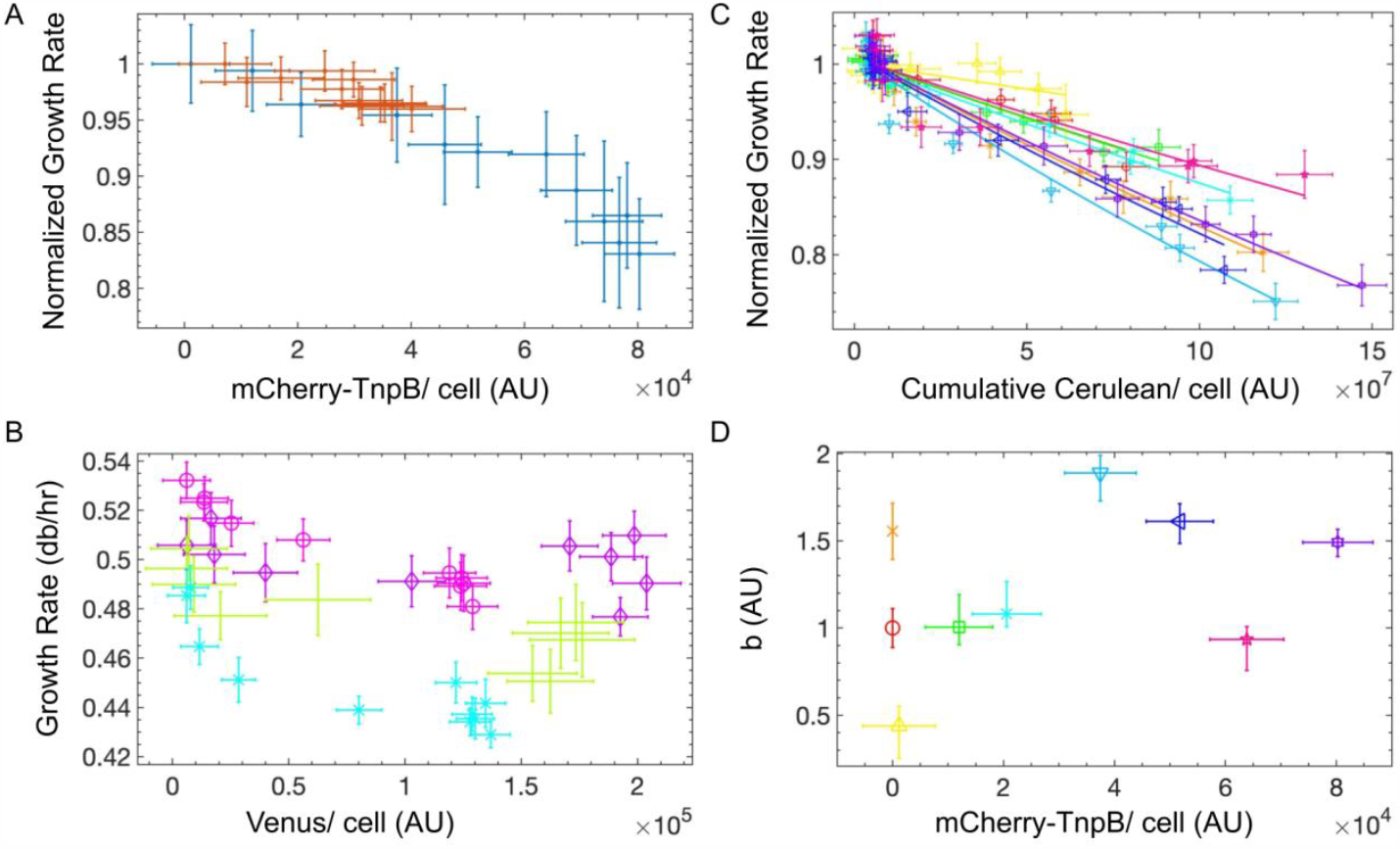
EFFECTS OF TNPB ON GROWTH. **Effects of TnpB on Bacterial Growth: (A)** TnpB expression in the presence of: (i) the active TE (blue) and (ii) a negative control plasmid without the active TE (red). Induction range is identical for both, [IPTG] = {0μM, 10μM, 20μM, 50μM, 100μM, 200μM, 2000μM}. However, the mCherry-TnpB response to induction is different as TnpB is unstable in the absence of other TE features (red). **(B)** Growth rate of immobilized strain without TnpB and no IPTG (purple), without TnpB and with IPTG (green), with TnpB and no IPTG (magenta), and with TnpB and with IPTG (light blue) versus Venus fluorescence per cell for all concentrations of aTc. Immobilized strains with the same concentrations of TnpA-Venus have a greater growth defect when TnpB is coexpressed compared to when only TnpA is expressed. **(C)** Growth rate is fit to an exponential decay function of Cumulative Cerulean determining *b* as described in **Supplementary Information SI.viii** for Tn4rev only (red, **◯**), TnpB with Tn4rev *in trans* for different IPTG concentrations [0μM (yellow, ∆), 10μM (green, □), 20μM (turquoise, ✳), 50μM (teal, ▽), 100μM (navy, ◁), 200μM (magenta, 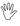) & 2000μM (violet, ✵)] and for TnpB *in cis* with Tn4rev (orange, X). **(D)** *b* increases with TnpB until it saturates. The outliers to this pattern correspond to the curves with R^2^ values < 0.9 (**Supplementary Table S6**). Data point for strain with TnpB introduced *in cis* to Tn4rev is plotted at zero point for mCherry-TnpB because it has multiple values along the fitted curve.

To determine the growth defect contributed by an excision event, we fit the growth rate for each strain to an exponential decay function of Cumulative Cerulean (titrated with [atc]) for each individual concentration of TnpB ([IPTG]) (**Fig. 3C**). In **Supplementary Table S6**, we list the R^2^ values of the fit of the exponential decay function for each curve. The coefficient of exponential decay (*b*) increases as a function of TnpB (**Fig. 3D, Supplementary Information SI.vii-viii**), suggesting that a combined effect of TnpA and TnpB is causing the defect. We suggest that TnpB co-expression increases the insertion efficiency of the TE. If the growth defect was simply the cumulative damage of TnpA and TnpB, the normalized growth curves would coincide as *b* would be consistent between them. We suggest that *b* is a measure of the successful insertion rate. As the insertion rate improves, the excision events that result in mutational damage increase, leading to greater growth decay.

### Modeling demonstrates that TnpB-improved insertion accounts for increased TnpA levels and TE activity

To understand how TnpB affects transposon dynamics, we analyzed a mean field theory in which the population averaged transposition rates, number of excised and unexcised plasmids and number of transposons are considered for a representative of the population (see **SII.i Supplementary Modeling** for details) which is evolved over time.

In our model, the representative starts with an experimentally determined average number of transposons, *X*_*o*_ = 12. The excision (*χ*, 0 ≤ *χ* ≤ 1) and insertion (*μ*) rates of each TE are proportional to the concentration of transposase molecules, which are generated by the TEs themselves. The single stranded transposon excises and then insert primarily into the lagging strand of the target during replication (0 ≤ *μ* ≤ 1) (*1, 5*) which results in a maintenance or loss of TE numbers and an average of at most half the daughter chromosomes containing a TE at the new site. We propose that TnpB extends the range of insertion rate, possibly doubling it by nicking the leading strand during insertion, resulting in the introduction of a TE in both daughter chromosomes. TnpB-assisted homing is considered by a rate of reintroduction of transposons to excision sites, *C*_*hom*_ (TE^-1^ [TnpB]^-1^ generation^-1^) ^4^. The final numbers and activity levels of the transposon in cells evolved (**Fig. 4A**) over experimental time scales and plotted in **Fig. 4B-E**. Our model recapitulates our experimental observations of an increase and earlier initiation of TnpA-Venus, total transposons, number of excision events and excised plasmids, with an increase in TnpB concentration.

**Figure 4:**
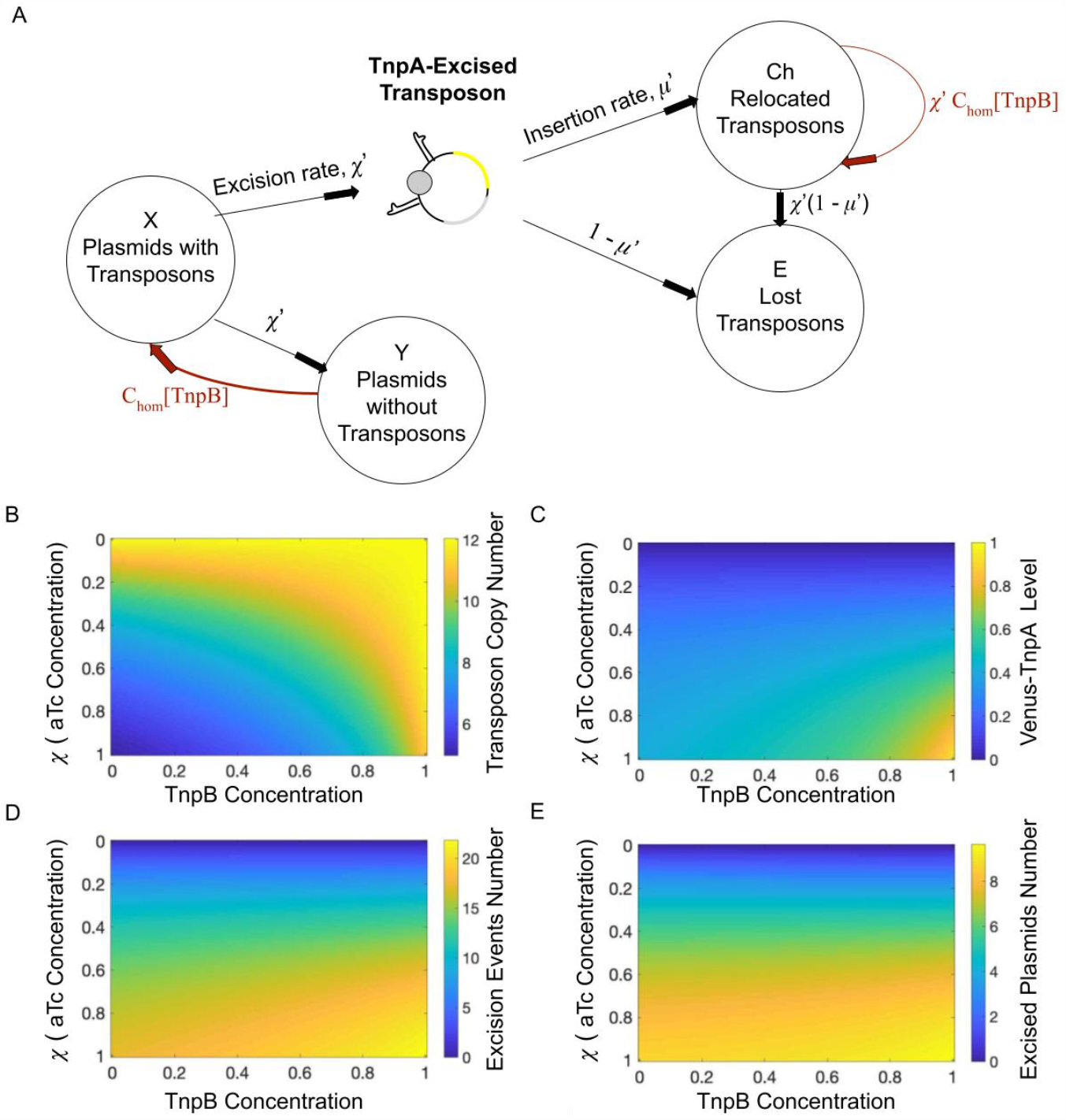
MODEL OF TRANSPOSON DYNAMICS. **Modeling of Transposon and TnpB Dynamics: (A)** Schematic of Mathematical Model: The rate equations described in **Supplementary SII.i** are evolved over time to generate the total number of transposons, number of plasmids with and without transposons in their original sites, the total number of excision events (Cerulean) and a measure of the relative transposase molecules (Venus). **(B-E)** The coupled differential equations were evolved over three generations to mimic experimental conditions and the above phase diagrams were generated. Phase diagrams were generated with the y-axis and x-axis measuring the induction quantities with aTc and IPTG respectively. Here, the induced excision rate per TE is a function of the aTc concentration as it induces TnpA, which executes excision. Tuning of the TnpB concentration from zero to one in the model accounts for increased peak values and earlier initiation of expression that we observe experimentally for **(B)** total transposon number (qPCR data), **(C)** transposase concentration (Venus-TnpA fluorescence data), **(D)** number of excision events (cumulative Cerulean fluorescence data) and **(E)** total excised plasmids (qPCR data).

## DISCUSSION

Inefficiencies in the transposition process can result in transposons being lost over time (**Fig. 2G, Supplementary Fig. S6, and Supplementary Information SI.vi**). This inefficiency compounded with the mutational damage of insertional sequence (IS) proliferation, makes IS extinction within hosts inevitable^41,42^. However, by mitigating the decay of TE numbers, TnpB can prolong TE lifetimes within hosts. We demonstrate that TnpB expression in conjunction with TnpA increases TE retention (**Fig. 2G**). TE retention further increases TnpA levels in cells (**Figs. 1B-D, and Supplementary Fig. S6)**, which induces greater excision numbers (**Figs. 2A-C,2H**). However, this increased TE retention comes at the cost of higher damage and resulting detrimental growth effects (**Figs. 3C-D**). Considering TnpB’s conservative function and abundance in bacterial and archaeal genomes, TnpB must have played an important role in the proliferation and maintenance of TEs throughout these genomes. Additionally, TnpB’s proliferation via IS200/IS605 transposons and positive impact on TE proliferation perhaps explain its abundance, shedding light on key dynamics leading to the evolutionary transition of the emergence of adaptive immunity in bacteria^2,34,35,43–45^.

We considered two mechanisms of TnpB interaction that allow it to increase TE retention: (1) TnpB allows an additional “homing” transposition mechanism that introduces a copy of the TE back into its original DNA excision site by cutting it and relying on homologous recombination to repair the cut^4^, or (2) TnpB increases the insertion efficiency of the TE into its TnpA-assisted transposition target site, reducing TE loss. Our qPCR results confirm that target site insertion increases with the introduction of TnpB. However, the increase in the number of plasmid-excision events with maximal TnpB vs. no TnpB induction (ratio of 1.67±0.22, **Fig. 2B**) is larger than the corresponding increase in the number of excised plasmids (ratio of 1.25±0.23, **Fig. 2E**), suggesting that every excision event does not result in a proportionate number of excised plasmids with a p-value of 0.028. This implies that TnpB is also aiding in homing of the TE back into its original site on the plasmid. Indeed, our model can only account for the large range of increase of excision events with increased TnpB only if we include TnpB-induced homing of the TE back into its original site on the plasmid (**Fig. 4D**). Importantly, increased insertional efficiency is required to account for the overall increase in the number of excised plasmids (**Fig. 4E**) and cannot be achieved with homing alone.

We find that co-expression of TnpA and TnpB results in growth defects, even in the absence of transposition. This suggests an interaction where the function of TnpB-RuvC may be analogous to RuvC’s function in Cas9 and Cas12. We propose that TnpA-TnpB may interact to collectively make dsDNA cuts and, within this mechanism, TnpB may nick the complementary strand of the TnpA-target site. This break would introduce the TE to both the target and its complementary strand, resulting in both daughter strands carrying TEs at this new site, doubling the maximum TE insertion rate. Simultaneously, TnpB on its own may make dsDNA cuts at the original excision site to execute TE homing^4^. Thus, we propose that TnpB has a dual effect on transposition—aiding in TnpA-executed transposition and providing an additional homing mechanism.

## ACKNOWLEDGEMENTS

This work was supported by the NSF Center for the Physics of Living Cells (PHY 1430124) and startup funds from the University of California.

## AUTHOR CONTRIBUTIONS

Conceptualization, T.E.K.; Methodology, D.K. and T.E.K.; Investigation, D.K., Formal Analysis, D.K.; Writing – Original Draft, D.K. and T.E.K.; Writing – Review and Editing, D.K. and T.E.K.; Funding Acquisition, T.E.K.; Resources, D.K. and T.E.K.; Supervision, T.E.K.

## METHODS

### Resource Availability

#### Lead Contact

Further information and requests for resources and reagents should be directed to and will be fulfilled by the lead contact, Thomas E. Kuhlman. All data reported in this paper will be shared by the lead contact upon request.

#### Data and Code Availability

Any additional information required to reanalyze the data reported in this paper is available from the lead contact upon request. Genetic sequences for the transposon constructs used in this study have been deposited with the NCBI GenBank under accession number OP581959, OP581957, OP581958, OP717084, and OP717085.

#### Materials Availability

Plasmids and strains used in this study are available from the lead contact upon request.

#### Strains and Media

Experiments were performed using *E. coli* K-12 MG1655 Δ*lac*^46–50^. Molecular cloning and plasmid manipulations were performed using *E. coli* NEB turbo as a host strain. Cells for measurement of the TE excision response function were grown in M63 minimal medium [100 mM KH2PO4, 15 mM (NH4)2SO4, 1 mM MgSO4, 1.7 μM FeSO4, 5e-5% (wt/vol) thiamine, pH adjusted to 7.0 with KOH] with 0.5% (vol/vol) glycerol as carbon source. Antibiotics were added to the medium as appropriate for plasmid maintenance, and different concentrations of aTc (Sigma Aldrich) and IPTG (ThermoFisher) were added to induce transposase and TnpB production respectively.

#### Plasmid Construction

The low copy number plasmid pJK14^51^ was used to host the TE in all experiments. pJK14 has a pSC101 replication origin. Plasmid copy number is tightly controlled through the positive feedback of the plasmid-encoded protein RepA^52^. Additionally, pJK14 is actively segregated to daughter cells through the pSC101 *par* system^53^.

Plasmid pJK14-Tn4rev was designed using Vector NTI software (Life Technologies) and synthesized *de novo* by GENEWIZ Gene Synthesis Services (GENEWIZ, Inc.).

Plasmid pZA31-*mCherry-TnpB* was designed using Vector NTI software (Life Technologies) and synthesized *de novo* by GENEWIZ Gene Synthesis Services (GENEWIZ, Inc.). The mCherry-TnpB cassette was inserted into pZA31 at the NheI-HF and XhoI cut sites.

***Construction of:***

1. **pJK14** The pJK14 fragment was amplified from the pJK14-Tn4rev with Phusion Flash Master Mix and the primers pJK14-I-SceI F : 5’-TAGGGATAACAGGGTAATGCATGCAAGCTCTAGACACGTGC-3’ pJK14-I-SceI R : 5’-ATTACCCTGTTATCCCTAGACGTCGGAATT-3’ The PCR product was then purified, digested with DpnI and I-SceI, religated, and transformed into NEB turbo. The construct was subsequently confirmed to have the correct size through digestion and gel electrophoresis. Absence of Tn4rev was confirmed using the below PCR primers and loss of Venus and Cerulean fluorescences. Pre-NheI-*tnpB* insert F: 5’-GGTCGTAGCAGTAGGATATTAAGACAAGAATTTAACCACTTAAAAACAAAACTA-3’ Pre-NheI-*tnpB* insert R: 5’-ACAATTGTTAGTATTAAATTGTGAGCGCTCACAATTATCAGCGAG-3’
2. **pJK14-Tn4rev-*mCherry*-*tnpB*** The mCherry-TnpB sequence was amplified from pZA31-mCherry-TnpB using the following primers: NheI-*RBS-mCherry* F: 5’-AAGCTAGCATTAAAGAGGAGAAAGGTACC-3’ NheI-*tnpB* R: 5’-AAAAAGCTAGCCTAACAAGTAGGTCTTACAAATTC-3’ The plasmid pJK14-Tn4rev and *mCherry-TnpB* cassette were both digested with NheI-HF, PCR-purified and ligated to generate pJK14-Tn4rev-mCherry-*tnpB*. The resulting plasmid was confirmed to have the correct size through digestion with I-SceI and the correct cassette orientation by digestion with KpnI-HF and subsequent gel electrophoresis. Additionally, the sequence was confirmed through PCR amplification with the below primers and Sanger sequencing at the Institute for Integrative Genome Biology at the University of California, Riverside. Pre-NheI-*tnpB* insert F: 5’-GGTCGTAGCAGTAGGATATTAAGACAAGAATTTAACCACTTAAAAACAAAACTA-3’ Pre-NheI-*tnpB* insert R: 5’-ACAATTGTTAGTATTAAATTGTGAGCGCTCACAATTATCAGCGAG-3’
3. **pJK14-Tn4rev-*tnpB*** The TnpB sequence was amplified from pZA31-*mCherry-tnpB* using the following primers: NheI-*RBS-tnpB* F: 5’-AAAAAGCTAGCATTAAAGAGGAGAAAGGTACCATGTTGATAACCTACAAACAAA-3’ NheI-*tnpB* R: 5’-AAAAAGCTAGCCTAACAAGTAGGTCTTACAAATTC-3’ The plasmid pJK14-Tn4rev and *tnpB* cassette were both digested with NheI-HF, pCR purified and ligated to generate pJK14-Tn4rev-*tnpB*. The resulting plasmid was confirmed to have the correct size through digestion with I-SceI and the correct cassette orientation by digestion with KpnI-HF and subsequent gel electrophoresis. Additionally, the sequence was confirmed through PCR amplification with the below primers and Sanger sequencing at the Institute for Integrative Genome Biology at the University of California, Riverside. Pre-NheI-*tnpB* insert F: 5’-GGTCGTAGCAGTAGGATATTAAGACAAGAATTTAACCACTTAAAAACAAAACTA-3’ Pre-NheI-TnpB insert R: 5’-ACAATTGTTAGTATTAAATTGTGAGCGCTCACAATTATCAGCGAG-3’
4. **pZA31-P**_***LlacOid***_**-SmR** The Spectinomycin resistance gene, *aadA*, was amplified from pTKRED using the following primers, NheI-P_*LlacOid*_-Smr F: 5’-ACGTCGCTAGCAATTGTGAGCGGATAACAATTGACAATTGTGAGCGCTCACAAGATACTGAGCACATCAGCAGGACGCACTGACCGAATTCATTAAAGAGGAGAAAGGTACCATGAGGGAAGCGGTGATCGC-3’ NheI-P_*LlacOid*_-Smr R: 5’-CACTCCTCGAGTTATTTGCCGACTACCTTGGTGATCTCGCCTTTCACGTAGTGGACA-3’ The resulting P_*LlacOid*_-Smr cassette and plasmid pZA31 were then digested with NheI and XhoI, PCR purified, ligated and screened on LB agar plates containing 100ug/ml spectinomycin and 2mM IPTG to generate pZA31-P_*LlacOid*_-SmR. The resulting plasmid was confirmed to be the correct size through digestion with SpeI-HF and gel electrophoresis. Additionally, the plasmid no longer conferred Cherry fluorescence. Insertion was confirmed through PCR using the following primer pair NheI-P_*LlacOid*_ F: 5’-ACGTCGCTAGCAATTGTGAGCGGATAACAATTGA-3’ NheI-P_*LlacOid*_-Smr R: 5’-CACTCCTCGAGTTATTTGCCGACTACCTTGGTGATCTCGCCTTTCACGTAGTGGACA-3’
5. **pJK14-P**_***Ltet-O1***_**-*venus*-*tnpA*** Plasmid pJK14-Tn4rev was used to create versions of the TE without the excision sites on each end. To construct the control plasmid expressing only Venus-TnpA (pJK14-*venus*-*tnpA*) the entire TE cassette was removed from pJK14-Tn4rev through digestion with I-SceI, dephosphorylation, and gel purification. The P_*Ltet-01*_-*venus*-*tnpA* fragment was amplified from pJK14-ISLEAD with Phusion Flash master mix and the primers, I-SceI-P_*Ltet-01*_*-venus* F: 5′-TTCCGACGTCTAGGGATAACAGGGTAATTTGACATCCCTATCAGTGATAGAGA-3′ *tnpA*-I-SceI R: 5′-GCTTGCATGCTAGGGATAACAGGGTAATTTATAGAGCTTTTGTTTGTAGGTTA-3′ The amplified fragment was digested with I-SceI and ligated into the pJK14-Tn4rev backbone to generate pJK14-*venus*-*tnpA*. To control expression of Venus-TnpA, pJK14-*tnpA* was transformed into and assayed with strain CZ071, the kind gift of Ido Golding, University of Illinois Urbana-Champaign, which is a K-12 MG1655 strain that constitutively expresses *tet* repressor.

#### Fluorescence Measurements

To measure the TE response functions (shown in **Figs. 2A-C, 3A,B,F,G**), MG1655 Δ*lac* cells from freezer-stock culture carrying the indicated version of the TE and pZA31 plasmid were grown overnight in LB + 25 μg/mL kanamycin + 34 μg/mL chloramphenicol. The optical density at 600 nm (OD600) of this culture was measured with a Bio-Rad SmartSpec Plus spectrophotometer, and an appropriate volume of the culture was added to 2.5-mL of M63 minimal medium + 0.5% (vol/vol) glycerol + 25 μg/mL kanamycin + 34 μg/mL chloramphenicol in a 20-mm glass test tube to yield a calculated initial OD600 = 0.0008. This tube was grown at 37 °C in a New Brunswick C76 water bath shaker with vigorous shaking until it reached an ∼OD600 = 0.15–0.20; at this OD, the cells are within the exponential phase of growth and have undergone ∼7-8 doublings. The culture was then diluted to OD600 = 0.01 in M63 minimal medium + 0.5% (vol/vol) glycerol + 25 μg/mL kanamycin + 34 μg/mL chloramphenicol and the appropriate concentration of IPTG (0uM, 10uM, 20uM, 50uM, 100uM, 200uM or 2000 uM) was added. The culture was then partitioned into 0.2mL aliquots for each well of a Corning Costar 96-well microplate (Clear, Round well, flat bottom: Well volume: 360uL, Cell growth area: 0.32cm2, TC-Treated, Sterile, Individually wrapped, 3596). Each plate consisted of 8 replicate wells of 12 different aTc concentrations (0ng/mL, 1, 2ng/mL, 3ng/mL, 5ng/mL, 9ng/mL, 20ng/mL, 30ng/mL, 50ng/mL, 80ng/mL, 100ng/mL and 200ng/mL). aTc and IPTG titrated the cells with transposase and TnpB inducers respectively. The plate was loaded into a BioRad Clariostar plate reader for measurements The cultures were maintained at 37 °C in the temperature-controlled environmental chamber of the plate reader with shaking at 300rpm. Optical density measurements and readings in three fluorescent channels, mCherry, Venus, and mCerulean3, were made every 10 minutes. Fluorescent excitation measurements were made at 561 nm (mCherry excitation, mCherry-TnpB), 514 nm (Venus excitation, Venus-TnpA levels), and 457 nm (mCerulean3 excitation, excision reporting), in that order.

The population-average fluorescence per cell was determined for each well of each experiment by applying a linear fit to the fluorescence vs. the optical density in the exponential phase of growth between ∼OD600 = 0.02 to 0.2, between generation 2 to 5, for mCherry, Venus, and mCerulean3. Fluorescence values for each combination of aTc and IPTG concentration combination were averaged over all replicate experiments (2-3 experiments per inducer combination with 8 replicates per experiment.)

#### Quantitative-PCR Protocol

qPCR was used to determine relative plasmid copy number, unexcised plasmid-transposon number and total transposon number for cells induced with varying amounts of aTc and IPTG inducer concentrations. qPCR was performed using a Bio-Rad CFX96 Touch Real-Time PCR thermal cycler with SsoAdvanced Universal SYBR Green Supermix (Bio-Rad). All four primer pairs listed in **Supplementary Table S7** were designed to have similar melting temperature values so that the successful annealing temperatures ranges of all reactions overlap. All reactions were performed concurrently on DNA templates from the same batches for uniform concentrations across all qPCR reactions. MG1655 Δ*lac* pJK14-P_*Ltet-01*_-Tn4rev pZA31-P_*LlacOid*_*-mCherry-tnpB* were grown overnight in LB + 25 μg/mL kanamycin + 34 μg/mL chloramphenicol. The optical density at 600 nm (OD600) of this culture was measured with a Bio-Rad SmartSpec Plus spectrophotometer, and an appropriate volume of the culture was added to 2.5-mL of M63 minimal medium + 0.5% (vol/vol) glycerol + 25 μg/mL kanamycin + 34 μg/mL chloramphenicol in a 20-mm glass test tube to yield a calculated initial OD600 = 0.008. The appropriate concentrations of aTc (0 ng/mL or 200ng/mL) and IPTG (0uM or 2000 uM) were added. This tube was grown at 37°C in a New Brunswick C76 water bath shaker with vigorous shaking until it reached an ∼OD600 = 0.15–0.20; at this OD, the cells are within the exponential phase of growth and have undergone ∼4-5 doublings. From this culture, 0.5mL was centrifuged, the supernatant was discarded and resuspended in UltraPure DNase/RNase-Free Distilled Water (Invitrogen) water. This suspension was used to generate six concentrations of a 5× dilution series of cells for use in qPCR. Amplification reactions were performed in 96 wells, 4 replicates of each dilution level and primer pair per experiment, in 0.2 ml 8-Tube PCR Strips (Bio-Rad™, clear #TBS0201) with 0.2 ml Flat PCR Tube 8-Cap Strips (Bio-Rad™, optical, ultraclear, #TCS0803). Optimum amplification conditions were determined for each by amplifying six concentrations of a 5× dilution series of MG1655 pJK14-Tn4rev pZA31-*mCherry-tnpB* using two-step amplification with a thermal gradient of 55–75°C. The optimum reaction conditions were determined to be: <u>Annealing temperature:</u> 59°C and <u>Extension time:</u> 30 seconds and used for all 4 reactions.

Primer pair (i) for number of cells: MG1655-nth F and MG1655-nth R (**Supplementary Table S7**) generates a 196bp amplicon from the nth gene of MG1655 under the optimum conditions with Tm = 86.0°C and efficiency ε = 93.74 ± 1.25% [defined by, where n is cycle number]. Primer pair (ii) for number of plasmids: DK-pJK14-qPCR F and DK-pJK14-qPCR R (**Supplementary Table S7**) generates a 367bp amplicon from the pJK14-PLtet-01-Tn4rev plasmid with Tm = 82.0°C and efficiency ε = 96.00 ± 0.76%. Primer pair (iii) for number of unexcised plasmids: pJK14-RE\IP-qPCR F and pJK14-RE\IP-qPCR R (**Supplementary Table S7**) generates a 226 bp amplicon from pJK14-PLtet-01-Tn4rev with Tm = 81.0°C and efficiency ε = 95.23 ± 0.66%. Primer pair (iv) for total number of transposons: tnpA-venus-qPCR F and tnpA-venus-qPCR R (**Supplementary Table S7**) generates a 148 bp amplicon from pJK14-Tn4rev with Tm = 87.0°C and efficiency ε = 99.15 ± 0.56%.

#### Statistical Analysis

Statistical analysis and error calculation of qPCR data is described in sections **SIII.iii. Quantitative-PCR Analysis** and **SIII.iv. Error Calculation for Quantitative-PCR** of the Supplementary Information respectively. Two-sided P-values listed in **Tables S4 and S5** were calculated by conducting a Student’s T-Test. For the fluorescence data, standard error or the mean was determined for data for each fluorescence color and induction level of aTc and IPTG. The 95% confidence intervals for the fitted parameters were extracted from the Hill function fit to the induction curves (**Figs. 1B,S2,2A,S4,S3A-B,S1B-C**) and used to determine the standard error of the mean for data in **Figs. 1D-F,2C-E,S3C-E**.

## Supplementary Information

### SI. SUPPLEMENTARY ANALYSIS

#### SI.i. Introduction of TnpB to the Inducible Transposon

TnpB is translationally fused to mCherry and introduced to the transposon *in trans* and individually induced with IPTG (0 μM, 10 μM, 20 μM, 50 μM, 100 μM, 200 μM & 2000 μM). Levels of mCherry-TnpB as measured by mCherry fluorescence increase upon titration with IPTG (**Supplementary Fig. S1B**). The data are fit to a Hill function of the form

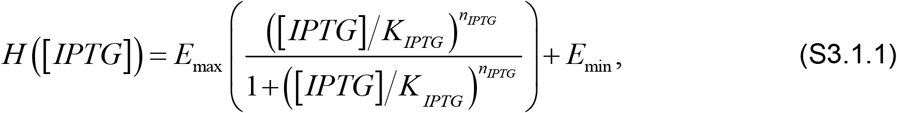

as described in the Methods section **Quantification of Fluorescence Per Cell**. Here, *E*_*max*_ is the overall scaling factor, *E*_*min*_ is the minimum value of the fit, *n*_*IPTG*_ is the Hill coefficient and *K*_*IPTG*_ is the LacI-IPTG dissociation constant. We find that *K*_*IPTG*_ *=* 61.53 μM and *n*_*IPTG*_ = 1.03.

Similarly, when TnpB is translationally fused to mCherry and introduced to the transposon *in cis* it is induced with aTc (0 ng/mL, 1 ng/mL, 2 ng/mL, 3 ng/mL, 5 ng/mL, 9 ng/mL, 20 ng/mL, 30 ng/mL, 50 ng/mL, 80 ng/mL, 100 ng/mL and 200 ng/mL). Levels of mCherry-TnpB as measured by mCherry fluorescence increase upon titration with aTc (**Supplementary Fig. S1C**). The data are fit to a Hill function of the form

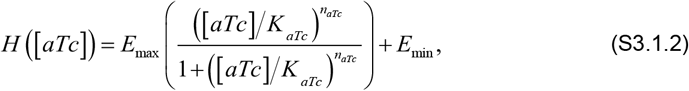

where parameters are defined similar to those in eq. (S3.1.1). We find that *K*_*aTc*_ *=* 285.77 ng/mL and *n*_*aTc*_ = 0.98.

#### SI.ii. Quantification of Venus-Transposase Concentration Per Cell

Venus-TnpA levels increase as a function of aTc concentration for all strains (**Figs. 1B-C and Supplementary Fig. S2**). The increase in Venus-TnpA levels for active TE strains (**Fig. 1B and Supplementary Fig. S2**) is more gradual than for the immobilized control strain with and without TnpB induction (**Fig. 1C**), which reach their saturation values past 20 ng/mL of aTc. Hence, excision alters the response and fluorescence profile of the transposon constructs. Venus-TnpA levels for all concentrations of aTc coincide for the strain containing only Tn4rev and the strain with TnpB introduced *in trans* with Tn4rev but is not induced with IPTG (**Supplementary Fig. S2**). Introduction of TnpB *in cis* with Tn4rev results in mCherry-TnpB increasing as a function of [aTc]. We note higher Venus-TnpA levels once expression of the P_*Ltet-01*_ promoter initiates (≥ aTc= 2ng/mL) for the case when TnpB is introduced *in cis* with Tn4rev relative to when we have Tn4rev only or when TnpB is introduced *in trans* with Tn4rev but is uninduced (0 μM IPTG). Furthermore, this difference in Venus-TnpA level between these cases increases with [aTc], as the concentration of TnpB increases (**Supplementary Fig. S2**). When TnpB is introduced *in trans* with Tn4rev, we can titrate the TnpB concentration individually with IPTG and find that this results in a gradual increase in Venus-TnpA with TnpB for each aTc concentration (**Figs. 1B, 1D-F**).

#### SI.iii. Quantification of Excision Rate Response Per Cell

In previous work^1^ we tracked Tn4rev excision events in individual cells and determined that with each excision event, we would observe a spike in Cerulean fluorescence in the cell which would then decay to zero. Thus, when we measure a population-averaged Cerulean fluorescence per cell, we obtain a measure of the active excisions. Hence, Cerulean fluorescence measures the population-averaged per cell rate of TE excision from plasmids. The excision rate increases as a function of aTc concentration for all three strains (**Supplementary Fig. S3A**). Excision rates for all concentrations of aTc for the strain containing only Tn4rev and the strain with TnpB introduced *in trans* with Tn4rev, with 0μM IPTG, are within error of each other (**Supplementary Fig. S3A**). Introduction of TnpB *in cis* with Tn4rev results in mCherry-TnpB increasing as a function of aTc. We note higher excision rates for the case when TnpB is introduced *in cis* with Tn4rev relative to the Tn4rev only and *in trans*, 0μM IPTG cases once excision initiates at aTc = 3ng/mL (**Supplementary Fig. S3A**). Titration of TnpB with IPTG for the strain with TnpB introduced *in trans* with Tn4rev results in an increase in excision rate of TEs per cell until the excision rate saturates at a maximum value (**Supplementary Figs. S3B,C**). With increased induction of TnpB with IPTG, we also observe earlier initiation of Cerulean fluorescence for lower concentrations of induction with aTc (**Supplementary Fig. S3D**).

#### SI.iv. Quantification of Excision Events Per Cell

Cumulative Cerulean fluorescence measures the population-averaged number of total excision events from plasmids per cell (see Methods section **Calculation of Cumulative Cerulean Fluorescence Per Cell**). The total number of excision events increases as a function of aTc concentration for all three strains (**Supplementary Fig. S4**). Introduction of TnpB *in cis* with Tn4rev results in mCherry-TnpB increasing as a function of aTc. We observe higher excision numbers for the case when TnpB is introduced *in cis* with Tn4rev relative to the Tn4rev only and *in trans*, 0μM IPTG cases once excision initiates at [aTc] = 3ng/mL - 4ng/mL (**Supplementary Fig. S4**). Titration of TnpB with IPTG for the strain with TnpB introduced *in trans* with Tn4rev results in a gradual increase in plasmid-excision events of TEs per cell for each aTc concentration (**Supplementary Fig. 2A,C**). With increased induction of TnpB with IPTG, we also observe earlier initiation of Cumulative Cerulean fluorescence for lower concentrations of induction with aTc (**Supplementary Fig. 2D**) and increased sensitivity to aTc (**Supplementary Fig. 2E**).

#### SI.v. Excision Response as a Function of Transposase Concentration

The Venus-TnpA concentration, the excision rate of TEs from plasmids, and the number of plasmid-excision events all increase with TnpB. Excision of the TE happens in response to the TnpA concentration in cells. Hence, we quantify Cerulean fluorescence as a function of Venus-TnpA concentration for all concentrations of TnpB (**Supplementary Fig. S5**). The data collapse to a single curve. As the concentration of transposase increases, the rate of excision events increases until it reaches its peak value.

#### SI.vi. Decay of Transposase numbers as a Function of TnpA and TnpB Concentrations

Guided by our qPCR results, we measured the decay in Venus-TnpA numbers per cell which is expected in response to a decay in TE numbers. The decay coefficients of the transposase concentration over time, proportionate to *β*, are determined for the strain with TnpB introduced *in trans* with Tn4rev for all concentrations of TnpA (aTc) and TnpB (IPTG). To determine *β* we fit the ratio of the fluorescence, *F*(*t*), to the optical density, *OD*(*t*), in the tail end of the exponential phase to an equation of the form,

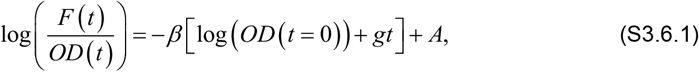

where *g* is the growth rate of the cells (determined as described in Methods), *t* is time in seconds, and *A* is a constant. Here, the fluorescence per optical density, ℱ_*C*_(t), decays as a function of time as follows:

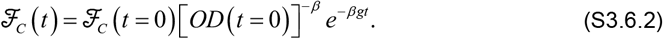

These exponent values, *βg*, are then normalized to values for the negative control strain and plotted (**Supplementary Fig. S6**). For lower concentrations of both proteins, the number of transposase molecules decays at a higher rate over time. Increase in TnpA and TnpB both result in maintenance of transposase concentration due to a lower decay rate over time. Further, TnpA and TnpB have a compounding effect on maintenance of TEs.

#### SI.vii. TnpB Increases Growth Rate Defect from Transposon Excision Events

Growth rate is fit to an exponential decay function of cumulative Cerulean of the form of **eq. S1.7.1** for Tn4rev only, TnpB with Tn4rev *in trans* for different concentrations of IPTG (0μM, 10μM, 20μM, 50μM, 100μM, 200μM & 2000μM) and for TnpB with Tn4rev *in cis* (**Fig. 3C**):

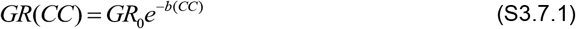

where GR is the population average growth rate of the cells, GR_0_ is population average growth rate of the cells with 0 ng/mL aTc, CC is the corresponding cumulative Cerulean fluorescence (CC) for each combination of [aTc] and [IPTG] and *b* is the coefficient of exponential decay. The coefficients of exponential decay (*b*) are plotted as a function of mCherry-TnpB in **Fig. 3D**. In **Supplementary Table S6**, we list the R^2^ values of the fit of the exponential decay function for each curve. We suggest that *b* is a measure of the successful insertion rate. As the insertion rate improves, the excision events that result in mutational damage increase, leading to the greater growth decay.

We note that, data for TnpB-Tn4rev *in trans* with 200μM IPTG (**Fig. 3C** magenta, 6.3843 × 10^4^ (AU) of mCherry-TnpB) were an outlier for the exponential decay fit (**Supplementary Table S6, Fig. 3D**). The growth rate initially drops with increase in excision events, but then remains consistently around 90% of its maximum value for higher values of cumulative Cerulean fluorescence. Data for this curve was a consistent outlier with each point on the curve being an average of 24 replicates. Curiously, Data for Tn4rev only case (**Fig. 3C** red) do not coincide with data for the TnpB-Tn4rev *in trans* with 0μM IPTG (**Fig. 3C** yellow, 1.1442 × 10^3^ (AU) of mCherry-TnpB) case. Data for Tn4rev only case instead coincides with data for TnpB-Tn4rev *in trans* with 10μM IPTG [**Fig. 3C** green, 1.1985 × 10^4^ (AU) of mCherry-TnpB]. These two outliers could be due to secondary functions of TnpB that we have not considered and are beyond the scope of our data.

#### SI.viii. Exponential Growth Defect Arises as a Direct Consequence of Genomic Integration

The observed exponential decay in normalized growth rate can be explained by a simple model where we consider the effect that integrations will have by disrupting essential chromosomal genes and thus cell viability. In the simplest model of this kind, we consider that there are two sub-populations of cells: those that grow normally, and those with transposon integrations disrupting all growth. In this binary model, there are *L* transcripts, each with a probability *w* of integrating and disrupting growth per generation, and the probability *q* of a cell having no integrations affecting growth per generation given by a binomial distribution evaluated at zero:

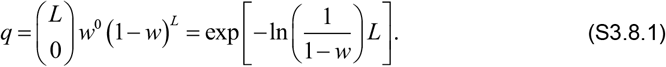

In our growth experiments, an exponentially growing individual cell, in the absence of integrations, will produce *g*_0_*dt* new individuals, in a time interval *dt*. This leads to a simple model of exponential growth of the form 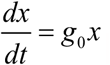. If we consider a binary model with a population *x* of normal cells and a population *y* of cells with no growth due to integrations, an individual of *x* will still produce *g*_0_*dt* new individuals but only a fraction *q* of these will be able to grow. This leads to the population model:

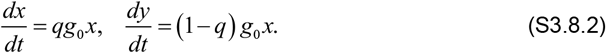

The total population of cells in this model grows as 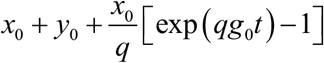. Thus, the measured growth rate would be *qg*_0_ and the normalized growth rate is just *q*. We fit eq. (S3.8.1) to the form exp[−*bL*] and make the identification *b* = −ln[1− *w*], which means *b* ≈ *w* for *w* ≪ 1. That is, *b* is approximately equal to the probability of a transposon transcript integrating and disrupting growth. Moreover, we expect that the rate of obtaining integrations affecting growth, *w*, is proportional to the overall rate of integration, *μ*. Consequently, this simple binary model recapitulates the exponential dependence of the growth rate on the number of transcripts and demonstrates that the exponential dependence implies that the growth defect, *b*, is expected to be directly coupled to the integration rate, *μ*.

More complex models of the impact of transposable element integration can be developed, with more than two sub-populations and more nuanced assumptions about the physiological effects. But we find that the dynamics of these models reduce to that of the two-rate model presented above, with renormalized parameters. An example of one such model is as follows. Let the number of cells with no chromosomal integrations harming their growth be *N*_0_, the population of cells with one integrant be *N*_1_, and so forth. Then a set of differential equations describing the population dynamics in exponential growth with growth rate *g*_0_ is

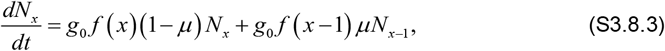

where *f* (*x*) is a monotonically decreasing function describing the inhibition of cell growth due to gene disruption by integrations, *μ* is the mutation rate, and the index *x* runs from 0 to some integer *x*_max_ such that the number of integrants is so high the cell cannot function and dies. Making the substitution (1− μ) = *q*, we have

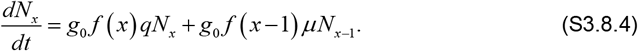

This is a lower triangular system of equations whose eigenvalues are the diagonals. After many generations, the largest eigenvalue will dominate and correspond approximately to the measured growth rate. Since *f* (*x*) is a monotonically decreasing function, this means the growth rate is *g*_0_ *f* (0)*q*. *f* (0) = 1 and, thus, the growth rate is *qg*_0_ and the normalized growth rate is *q*. This is the same result as the binary model discussed above.

### SII.i SUPPLEMENTARY MODELING

We consider a mean field theory in which the population averaged transposition rates, number of excised and unexcised plasmids and number of transposons are considered for a single representative of the population. This representative is evolved over time and plasmid and transposon numbers are tracked. The representative cell starts with a constant number of plasmids, *X*_o_, which each have a transposon. Hence the total initial number of transposons is *X*_o_. Each transposon has an inducible excision rate of *χ* per generation, where 0 ≤ *χ* ≤ 1. The effective excision rate is determined by

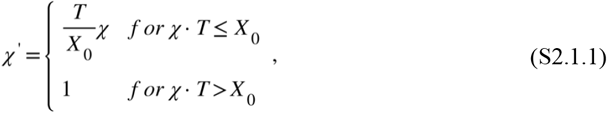

where *T* is the average total number of transposons per cell and *X*_o_ is a constant equal to the average total number of plasmids per cell which is the number of transposons per cell at t = 0. The effective transposition rates accounts for the level of transposase, which executes transposition, and is produced by the transposons themselves. The excision rate reaches a peak value beyond which an increase in transposase molecules does not further increase its value as we experimentally determine and show in **Supplementary Fig. S5**. *T* can change as the cell evolves. This is due to introducing a tunable successful insertion rate *μ*’ for each transposon once it has been excised and attempts to relocate. In the absence of TnpB, the transposons insert into the lagging strand of the target during replication. Thus, in the absence of TnpB, such that the insertion rate per generation *μ* satisfies 0 ≤ *μ* ≤ 1. We propose that TnpB extends the range of effective insertion rate by through an unknown mechanism such as nicking the leading strand, resulting in the introduction of a TE in both daughter chromosomes. This is consistent with the dynamic range here of *b* (**Fig. 3D**) and *βg* (**Supplementary Fig. S6**), the decay coefficient of the transposase molecules. Thus, in the presence of TnpB the insertion rate per TE becomes

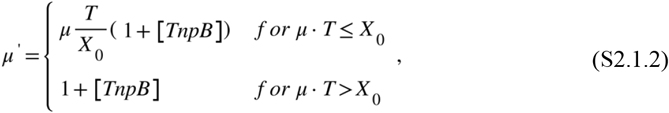

where [TnpB] is a measure of the concentration of TnpB, 0 ≤ [TnpB] ≤ 1 and [TnpB] = 1 corresponds to saturation of the cells with TnpB.

During each cycle, each transposon has a probability *χ*’ of excising per generation, leaving behind one empty plasmid strand, and one with the DNA complementary to the TE. These excised transposons then have a probability *μ*’ of successfully reinserting at a new location per generation. The schematic in **Fig.4A** provides an illustration of how the cell will evolve over time. The following coupled differential equations, along with the expressions above for *χ*’ and *μ*’, are concurrently solved numerically to track the changes in the transposition rates and the plasmid and transposon numbers over time t.

Number of plasmids with transposons *X*(*t*):

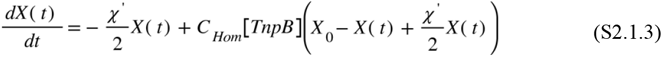

Number of reinserted transposons *Ch*(t):

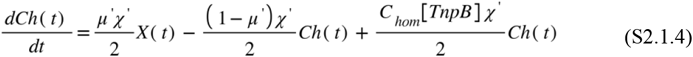

Number of plasmids without transposons *Y*(t):

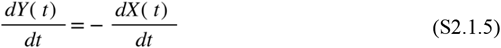

Total number of transposons, *T*(t):

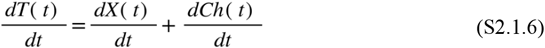

or

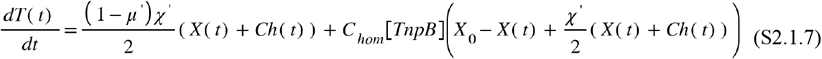

*C*_*hom*_ is the number rate of reintroduction of transposons reintroduced to excised plasmids sites (cell^-1^ [TnpB]^-1^ generation^-1^). For plasmid sites, the rate of transposon reintroduction through homing is proportional to the number of unexcised plasmids in the cell. However, for chromosomal sites, homing must occur within the same generation prior to cell division. Thus, this term is proportional to only the excisions that have just occurred. The cells are evolved over 3 generations using a forward Euler algorithm to mimic experimental timescales. The final numbers for each variable [**Equations (S2.1.3) – (S2.1.7)**] are plotted in **Figs. 4B-E**. It is clear from **Equation (S2.1.7)** that in the case when μ’ < 1 and *χ*’≠ 0, we have an exponential decay of transposons from transposition. When μ’ > 1 and *χ*’≠ 0, there is an exponential increase in transposons, which we do not experimentally observe, possibly due to suppression of the population average number of transposons per cell due to the damage of transposon proliferation.

### SIII. SUPPLEMENTARY METHODS AND STATISTICAL ANALYSIS

#### SIII.i. Quantification of Fluorescence Per Cell

The Venus, Cerulean and Cherry fluorescences per cell were determined as described for each combination of aTc and IPTG quantity for the strains: 1) Negative Control Strain for background subtraction, MG1655 pJK14 pZA31-P_*LlacOid*_-SmR, 2) Transposons Tn4rev only, MG1655 pJK14-P_*Ltet-01*_-Tn4rev pZA31-P_*LlacOid*_-SmR, 3) Trans-combination of Tn4rev and TnpB, MG1655 pJK14-P_*Ltet-01*_-Tn4rev pZA31-P_*LlacOid*_-*mCherry-tnpB*, 4) Cis-combination of Tn4rev and TnpB, MG1655 pJK14-P_*Ltet-01*_-Tn4rev-*mCherry-tnpB* pZA31-P_*LlacOid*_-SmR, and 5) Immobilized strain without TnpB, CZ071 pJK14-P_*Ltet-01*_-*tnpA* pZA31-P_*LlacOid*_-SmR and 6) Immobilized strain with TnpB, CZ071 pJK14-P_*Ltet-01*_-*tnpA* pZA31-P_*LlacOid*_-TnpB.

Background subtractions corresponding to the respective aTc and IPTG concentrations were subtracted from the fluorescence measurements to determine fluorescence per cell for each “color”. The resulting Venus and Cerulean fluorescence values per cell were plotted vs. aTc and fitted to a Hill function of the below form. The resulting mCherry fluorescence values per cell were plotted vs. IPTG and fitted to a Hill function of the below form.

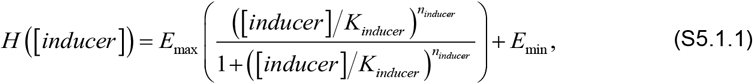

where *E*_*max*_ is the overall scaling factor, *E*_*min*_ is the minimum value of the fit, *n*_*inducer*_ is the Hill coefficient and *K*_*inducer*_ is the activation coefficient. The quantitative features of the responses are extracted. We use the inflection point to compare relative initiation points of the Hill fits. The inflection point is calculated using the following equation

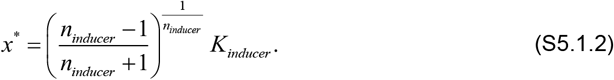

The inflection point corresponds to the following value of the Hill function,

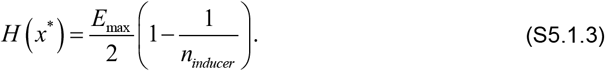

The slope at the inflection point is used as a measure of the sensitivity of the system to induction and is calculated using the equation

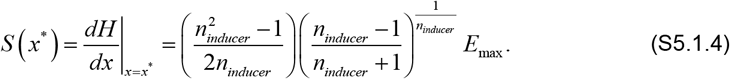

The error in the location of the inflection is determined using the 95% confidence interval of the Hill fit variables and the following equation

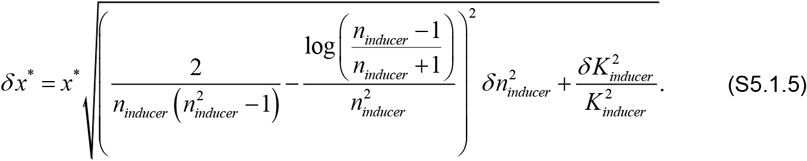

where *δn*_*inducer*_ is the standard error of the mean of the Hill coefficient. The error in the slope of the inflection is determined using the 95% confidence interval of the Hill fit variables and the following equation

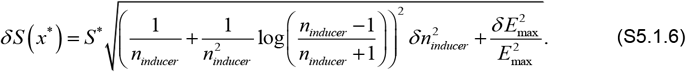

#### SIII.ii. Calculation of Cumulative Cerulean Fluorescence Per Cell

In previous work, we demonstrate that excision events are followed by spikes of Cerulean fluorescence. Thus, we measure excision rate per cell as the slope of Cerulean fluorescence vs. optical density, as described in the section **Quantification of Fluorescence per Cell**. In **Supplementary Fig. S7**, we illustrate the bimodal nature of the slope of excision rate that we observe during the exponential growth phase of *E. coli*. In **Supplementary Fig. S7**, we plot the initial excision rate, *m*_1_, starting at the lowest OD∼0.01 until the slope changes (green selection region of schematic) at point OD*. However, to determine the total number of excision events from the plasmids in the exponential phase, we integrate Cerulean fluorescence per cell (AU) over time. We measure *m*_2_ to make the following calculation of cumulative Cerulean or total plasmid excision events during exponential growth.

Cerulean Fluorescence per cell is the rate of excision from plasmids,

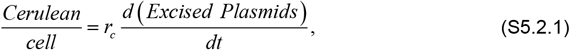

where, *r*_*c*_ is the fluorescence per plasmid-excision event. The integral of Cerulean Fluorescence per cell over time is the total number of excision events from plasmids:

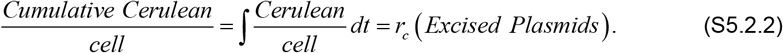

Executing the integral over OD, we obtain

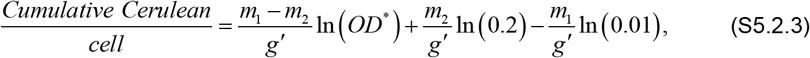

where *g*’= *g* ln(2) and *g* is the growth rate of the cells.

#### SIII.iii. Quantitative-PCR Analysis

Quantitative PCR reactions with all four primer pairs from **Supplementary Table S7** were performed concurrently on the dilution series of cells with reactions performed in quadruplicate. The quantity of cells is conserved between reactions for each dilution factor *d*_*i*_ (*d*_*i*_∈[5^1^,5^6^]) and the amplification from the MG1655 *nth* chromosomal locus located near the replication terminus^2–4^ allows for a measurement proportional to the number of cells. This allows the following calculation of the relative number of each gene per cell:

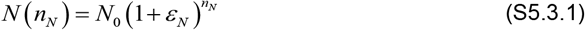

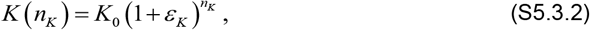

where *N* is the number of amplicons in the reaction, *ϵ*_*N*_ is the efficiency of the reaction, and *n*_*N*_ is the cycle number for qPCR reactions 2-4 (**Supplementary Table S7**): plasmid copy number, total transposon number, or unexcised plasmid number. Here, *K* is the number of MG1655 *nth* gene amplicons in the reaction (primer pair 1), *ϵ*_*K*_ is the efficiency of the reaction and *n*_*K*_ is the cycle number. When we consider the ratio of these equations at the threshold cycle number,

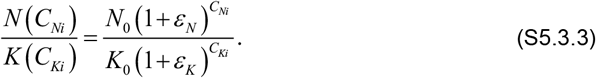

Using the same arbitrary threshold line for all reactions to determine the threshold cycle numbers of the *i*th sample (*C*_*Ni*_/*C*_*Ki*_, *i* = 1,2…6) makes the left-hand side of eq. (S5.3.3) equal to one and this gives us a relative value of the initial concentration of the sequence of interest per cell, i.e., *N*(*C*_*Ni*_) = *K*(*C*_*Ki*_), to obtain

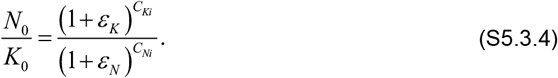

These relative values are determined for each combination of aTc (0 ng/mL and 200 ng/mL) and IPTG (0 mM and 2 mM) concentration for comparison.

#### SIII.iv. Error Calculation for Quantitative-PCR

Quantitative PCR reactions with all four primer pairs from **Supplementary Table S7** were performed concurrently on the dilution series of cells with reactions performed in quadruplicate. The average relative number of each amplicon per cell for each dilution level, *i* = 1, 2, …, 6, was determined from four replicate reactions:

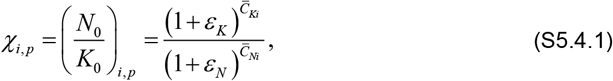

where *p* =1, …, *P*, is the experiment number, and 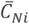 and 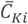 are the threshold cycle numbers of the *i*^th^ dilution level averaged over the four replicates. The standard deviation of the relative number of each amplicon per cell for each dilution level, *i* = 1, 2, …, 6, was determined as follows:

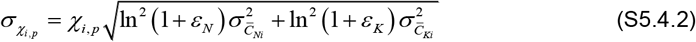

where 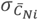 and 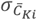 are the standard deviations of threshold cycle numbers of the four replicates of the *i*^th^ dilution level. The [χ_i,p_,σ_χi,p_] values from all experiments for each combination of aTc and IPTG inducer concentrations were then pooled. These pooled values were averaged over all experiments and dilution levels to determine the average amplicon number per cell for cells induced with a particular combination of IPTG and aTc concentration,

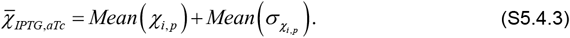

The standard deviation of the pooled χ_i,p_ values was determined to be σ_χ_. The total standard deviation of the amplicon number per cell for each IPTG and aTc concentration was calculated by accounting for the propagating error from σ_χi,p_, the values of which have a standard deviation of σ_δχi,p_. The total standard deviation of each 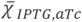 value is then

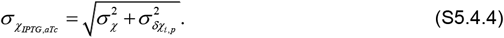

The uncertainties were determined by calculating the standard error of the mean:

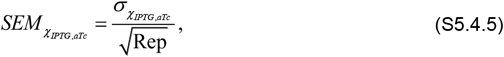

where Rep is the number of successful replicates.

## SUPPLEMENTAL FIGURES

**Figure S1:**
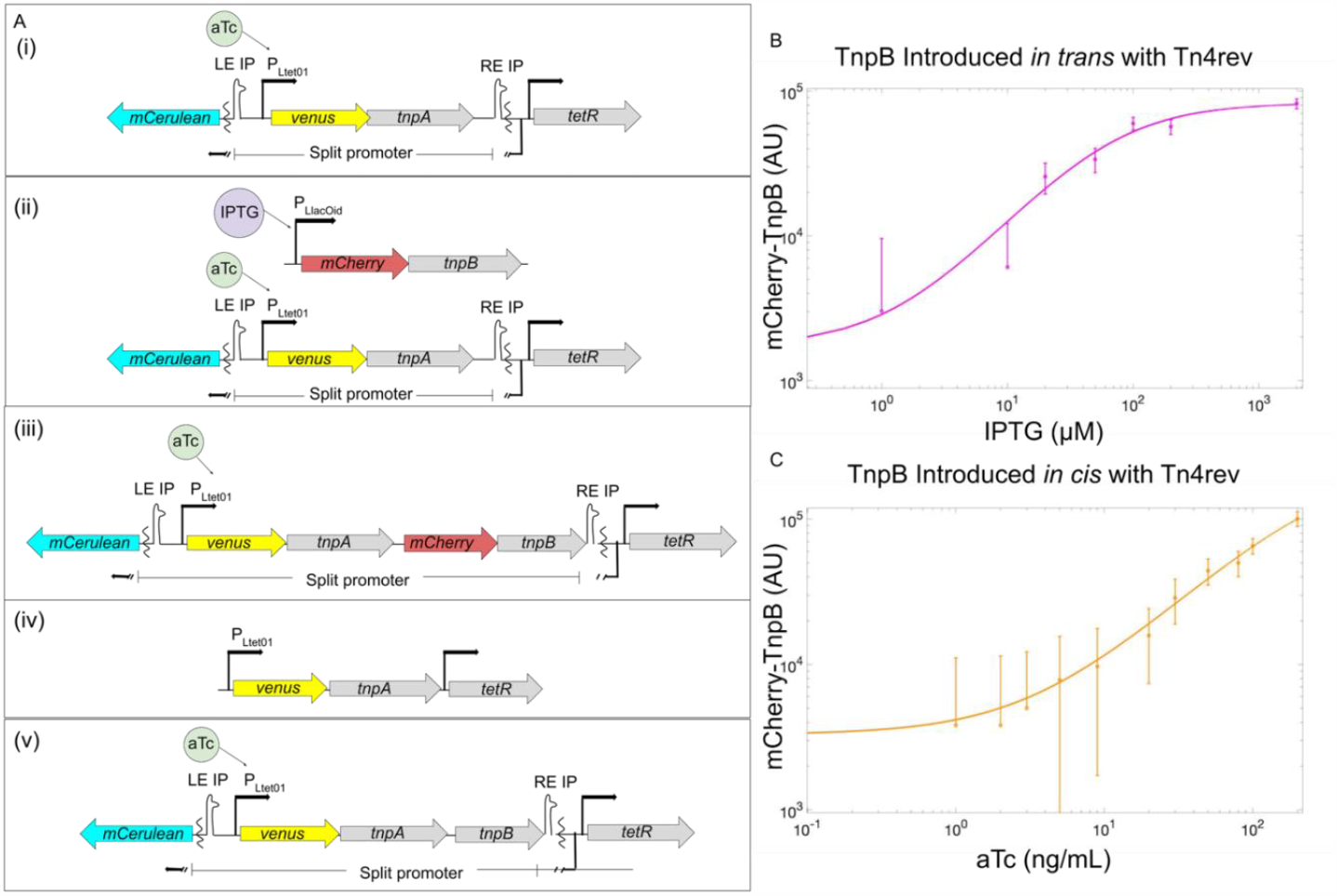
(A) Genetic Constructs for Inducible and Trackable Transposons: All transposons constructed for experiments. The strain which has Tn4rev only (i), MG1655 ∆*lac* pJK14-P_*Ltet-01*_-Tn4rev & pZA31-P_*LlacOid*_-SmR. TnpB introduced to the transposon *in trans* (ii), MG1655 ∆*lac* pJK14-*P*_*Ltet-01*_*-Tn4rev &* pZA31-P_*LlacOid*_-*mCherry-tnpB* and *in cis* (iii), MG1655 ∆*lac* pJK14-P_*Ltet-01*_-*tn4rev-mCherry-tnpB &* pZA31-P_*LlacOid*_-SmR. A version of the transposon (iv) without the excision sites, rendering it immobile. To the immobile transposon, we introduce, *in trans*, both TnpB: CZ071 pJK14-P_*Ltet-01*_-*tnpA &* pZA31-P_*LlacOid*_-*mCherry-tnpB*, and its negative control plasmid: CZ071 pJK14-P_*Ltet-01*_*-tnpA &* pZA31-P_*LlacOid*_-SmR. A control for the mCherry fusion to TnpB is also constructed (v), MG1655 ∆*lac* pJK14-*P*_*Ltet-01*_*-tn4rev-tnpB &* pZA31-P_*LlacOid*_-SmR. **(B)** TnpB is translationally fused to mCherry, introduced to the transposon *in trans* and individually induced with IPTG (0 μM, 10 μM, 20 μM, 50 μM, 100 μM, 200 μM & 2000 μM). **(C)** TnpB is translationally fused to mCherry and introduced to the transposon *in cis*. mCherry-TnpB and Venus-TnpA are induced simultaneously with aTc (0 ng/mL, 1 ng/mL, 2 ng/mL, 3 ng/mL, 5 ng/mL, 9 ng/mL, 20 ng/mL, 30 ng/mL, 50 ng/mL, 80 ng/mL, 100 ng/mL and 200 ng/mL).

**Figure S2:**
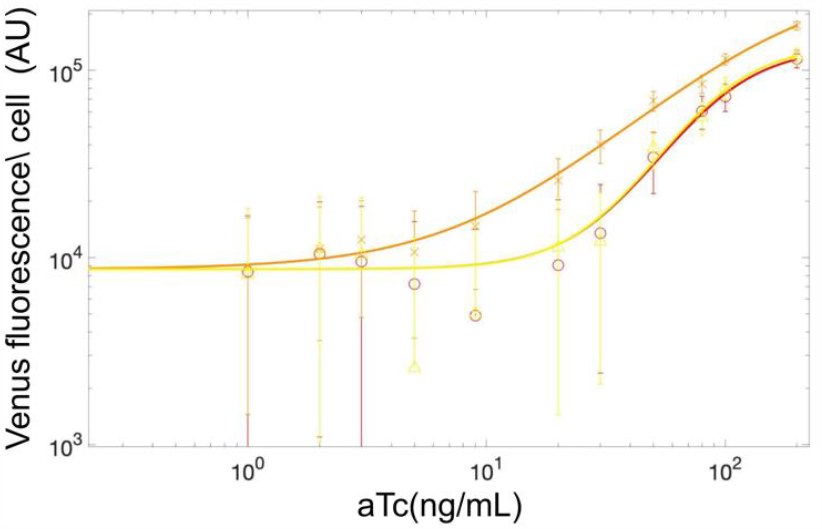
Quantification of Venus-Transposase Concentration to aTc Induction: Venus fluorescence data as a function of aTc concentration (0ng/mL, 1ng/mL, 2ng/mL, 3ng/mL, 5ng/mL, 9ng/mL, 20ng/mL, 30ng/mL, 50ng/mL, 80ng/mL, 100ng/mL and 200ng/mL) for **(A)** (i) Tn4rev only strain: MG1655 ∆*lac* pJK14-*P*_*Ltet-01*_-Tn4rev & pZA31-P_*LlacOid*_-SmR (red, **◯**), (ii) TnpB introduced *in trans* with Tn4rev at [IPTG] = 0: MG1655 ∆*lac* pJK14-P_*Ltet-01*_-Tn4rev & pZA31-P_*LlacOid*_*-mCherry-tnpB* (light green, ∆), (iii) TnpB introduced *in cis* with Tn4rev: MG1655 ∆*lac* pJK14-P_*Ltet-01*_-Tn4rev-*mCherry-tnpB &* pZA31-P_*LlacOid*_-SmR (orange, X).

**Figure S3:**
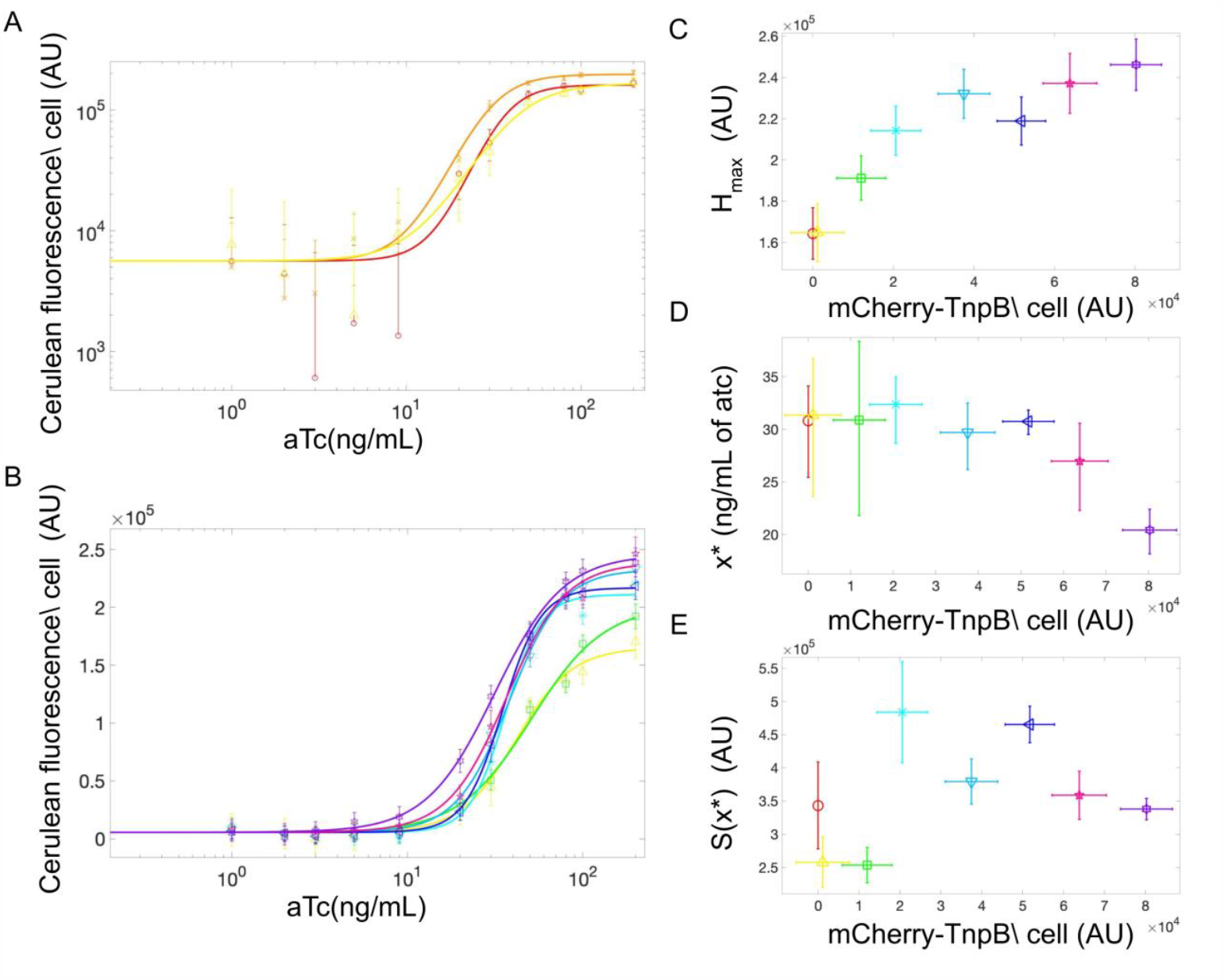
Titration of Excision Rate with Anhydrotetracycline (aTc): (**A,B)** Cerulean fluorescence data for [aTc] = {0ng/mL, 1ng/mL, 2ng/mL, 3ng/mL, 5ng/mL, 9ng/mL, 20ng/mL, 30ng/mL, 50ng/mL, 80ng/mL, 100ng/mL and 200ng/mL} for **(A)** (i) Tn4rev only strain: MG1655 ∆*lac* pJK14-P_*Ltet-01*_-Tn4rev & pZA31-P_*LlacOid*_-SmR (red, **◯**), (ii) TnpB introduced *in trans* with Tn4rev, [IPTG] = 0: MG1655 ∆*lac* pJK14-P_*Ltet-01*_-Tn4rev & pZA31-P_*LlacOid*_-*mCherry-tnpB* (light green, ∆), (iii) TnpB introduced *in cis* with Tn4rev: MG1655 ∆*lac* pJK14-P_*Ltet-01*_-01-Tn4rev-*mCherry-tnpB &* pZA31-P_*LlacOid*_-SmR (orange, X), and **(B)** TnpB is introduced *in trans* with Tn4rev, MG1655 ∆*lac* pJK14-P_*Ltet-01*_-Tn4rev pZA31-P_*LlacOid*_-*mCherry-tnpB* for different concentrations of IPTG [0μM (yellow, ∆), 10μM (green, □), 20μM (turquoise, ✳), 50μM (teal, ▽), 100μM (navy, ◁), 200μM (magenta, 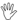) & 2000μM (violet, ✵)]. Cerulean fluorescence data sets are each fit to a Hill function (same color as data points for each) of the form of eq. **(S3.1.2)** and quantitative features of the responses are extracted and included in **C-E** and **Supplementary Table S2** for the strain with TnpB *in cis* with Tn4rev as it does not have a singular mCherry-TnpB value. **(C)** Higher Cerulean levels with TnpB induction when TnpA is maximally expressed (*H*_*max*_). **(D)** Earlier initiation of Cerulean expression (inflection point, *x\**) for lower aTc concentrations with increased TnpB induction. **(E)** Indiscernible pattern of excision sensitivity to aTc concentration (slope at inflection point *S(x*)*) with increased TnpB concentration.

**Figure S4:**
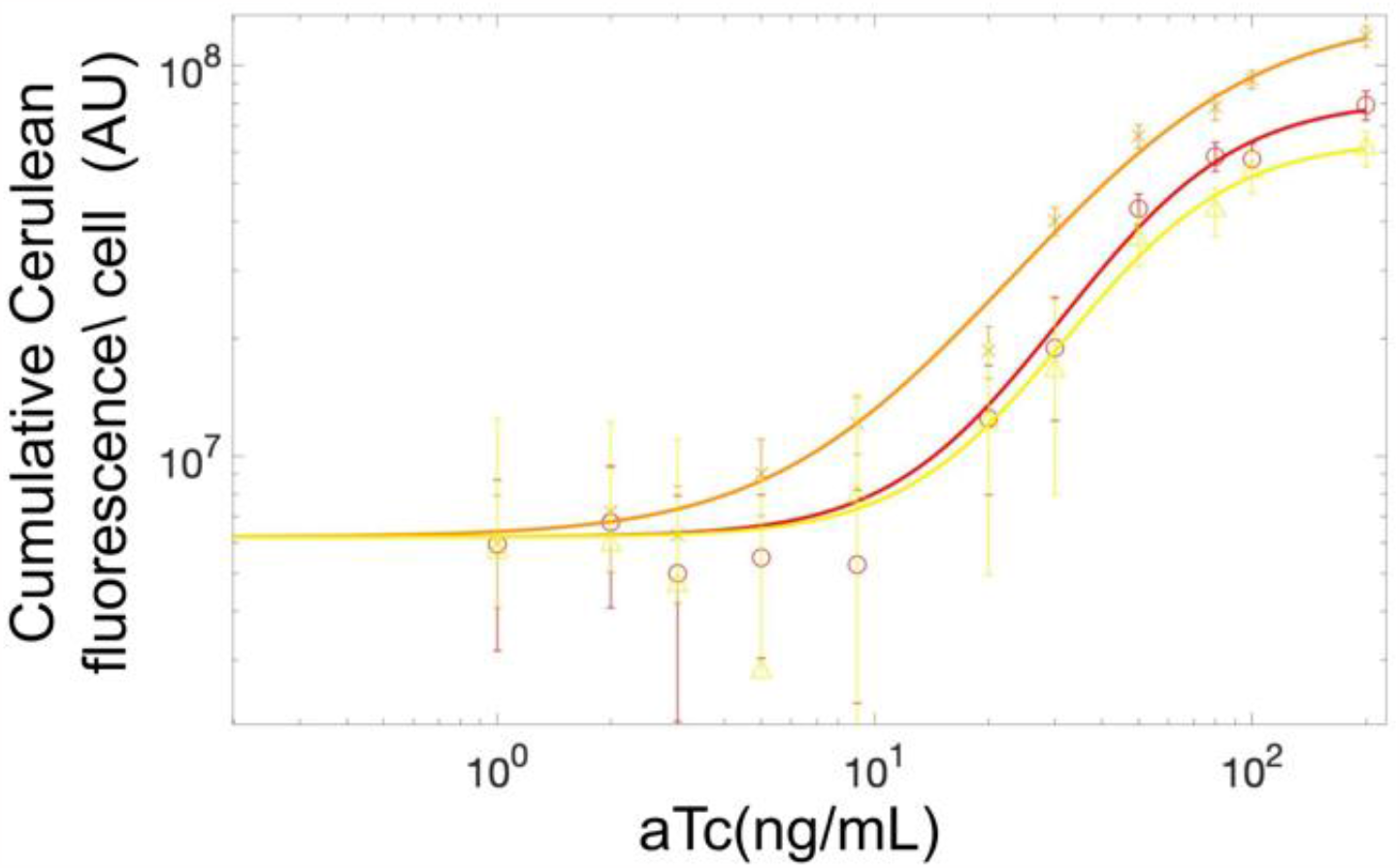
Titration of Excision Event Number with Anhydrotetracycline (aTc): Cumulative Cerulean fluorescence data as a function of inducer concentration for (0ng/mL, 1ng/mL, 2ng/mL, 3ng/mL, 5ng/mL, 9ng/mL, 20ng/mL, 30ng/mL, 50ng/mL, 80ng/mL, 100ng/mL and 200ng/mL) for (i) Tn4rev only strain: MG1655 ∆*lac* pJK14-P_*Ltet-01*_-Tn4rev & pZA31-P_*LlacOid*_-SmR (red, **◯**), (ii) TnpB introduced *in trans* with Tn4rev: MG1655 ∆*lac* pJK14-P_*Ltet-01*_-Tn4rev & pZA31-P_*LlacOid*_-*mCherry-tnpB* (light green, ∆), and (iii) TnpB introduced *in cis* with Tn4rev: MG1655 ∆*lac* pJK14-P_*Ltet-01*_-01-Tn4rev-*mCherry-tnpB &* pZA31-P_*LlacOid*_-SmR (orange, X).

**Figure S5:**
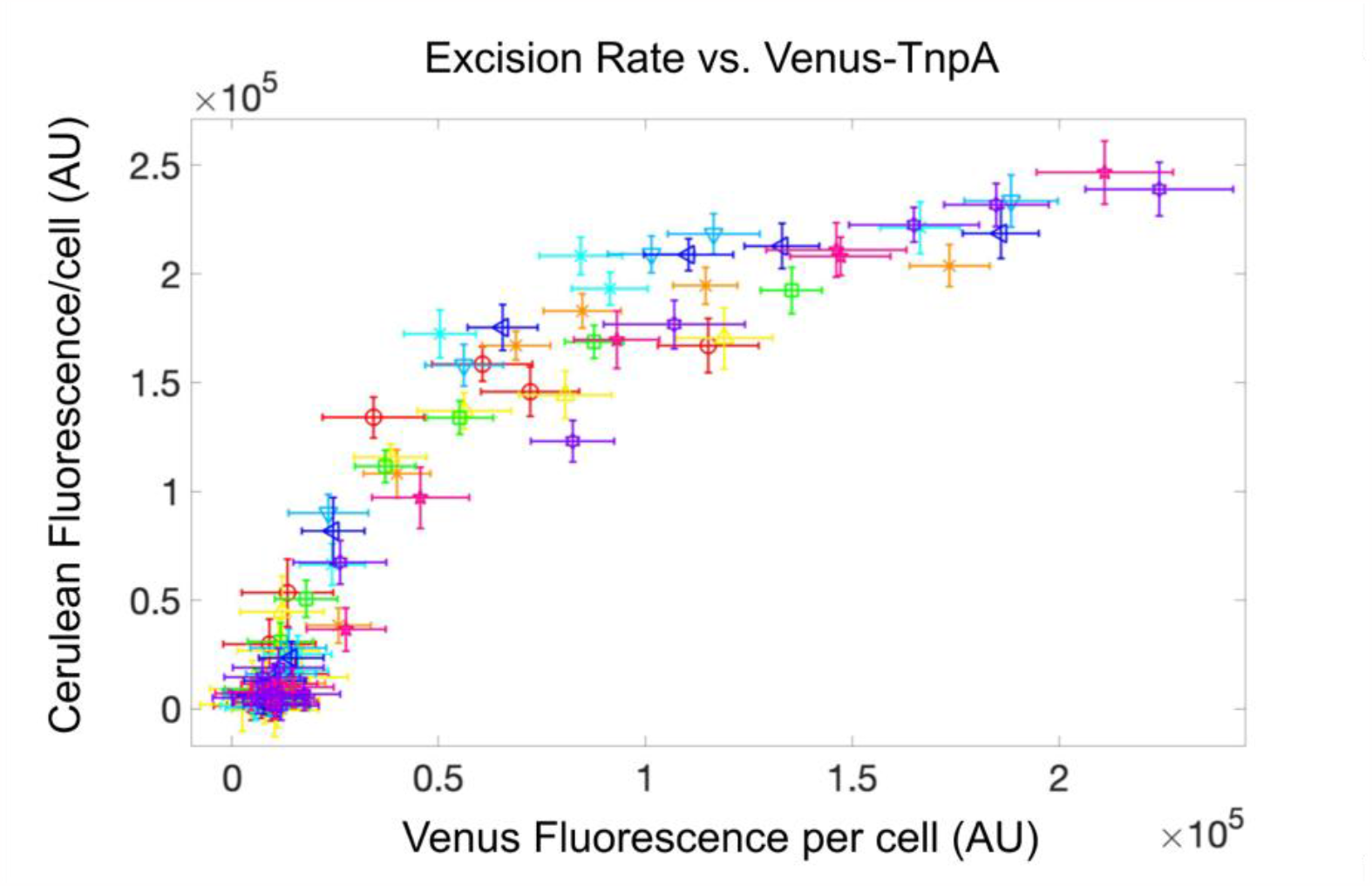
Excision Rate Increases and Peaks with Transposase Concentration: Excision rate (Cerulean fluorescence per cell) vs. transposase number (Venus fluorescence per cell) is plotted for: Tn4rev only strain: MG1655 ∆lac pJK14-PLtet-01-Tn4rev & pZA31-PLlacOid-SmR (red, ◯), TnpB introduced in trans with Tn4rev: MG1655 ∆lac pJK14-PLtet-01-Tn4rev pZA31-PLlacOid-mCherry-tnpB for different concentrations of IPTG (0μM (yellow, ∆), 10μM (green, □), 20μM (turquoise, ✳), 50μM (teal, ▽), 100μM (navy, ◁), 200μM (magenta, 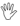) & 2000μM (violet, ✵)), and TnpB introduced in cis with Tn4rev: MG1655 ∆lac pJK14-PLtet-01-Tn4rev-mCherry-tnpB & pZA31-PLlacOid-SmR (orange, X). Both Venus-TnpA concentration and the excision rate of TEs from plasmids increase with TnpB. As the concentration of transposase increases, the rate of excision events increases proportionally until it reaches its peak value.

**Figure S6:**
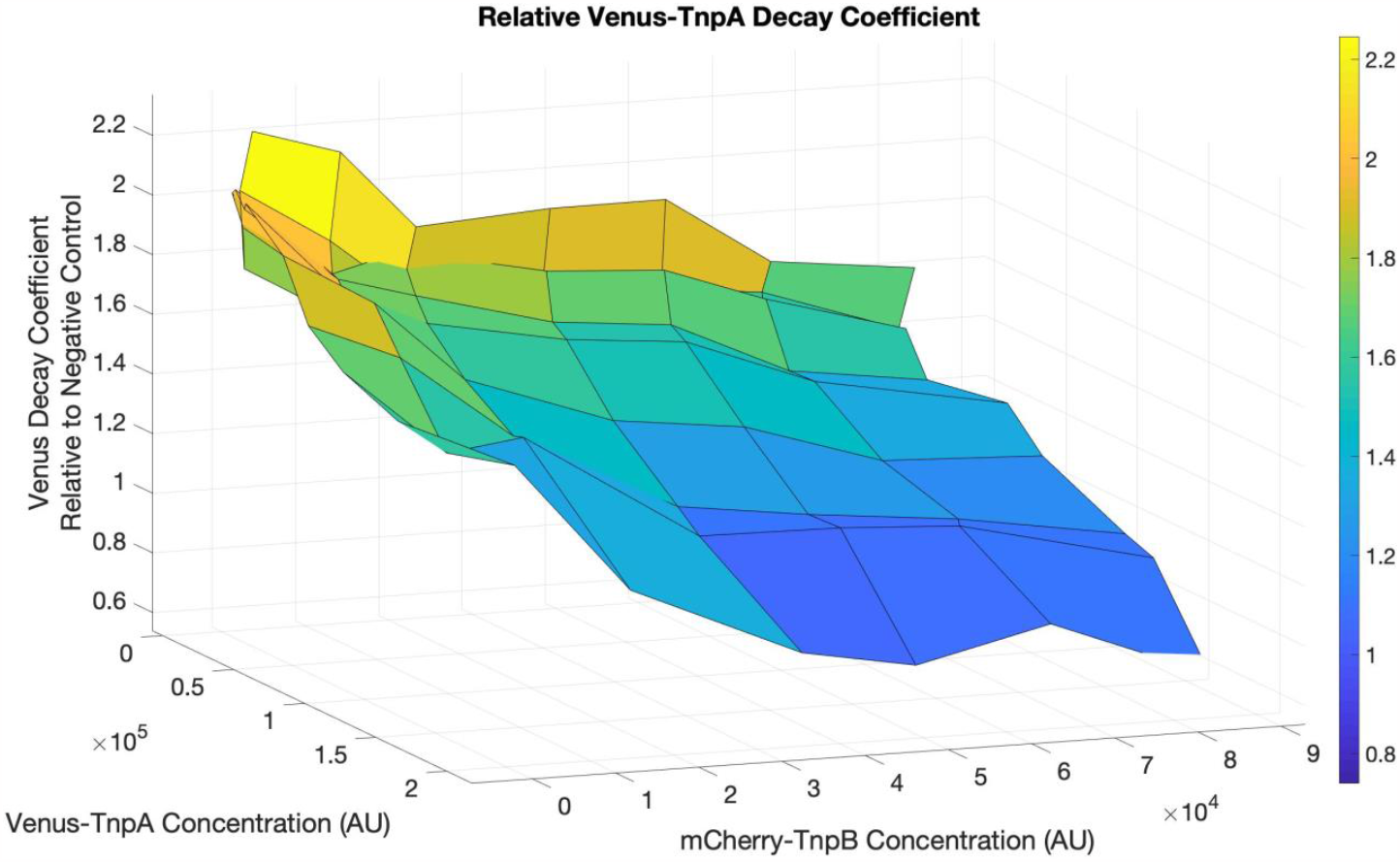
Decay of Number of Transpose Molecules per cell over time: The decay coefficients of the transposase concentration over time, *βg*, for the strain with introduction of TnpB *in trans* with Tn4rev are normalized to values for the negative control strain and plotted for all concentrations of TnpA and TnpB are plotted. For lower concentrations of both proteins, the number of transposase molecules decays at a higher rate over time. Increase in TnpA and TnpB both result in maintenance of transposase concentration due to a lower decay rate over time.

**Figure S7:**
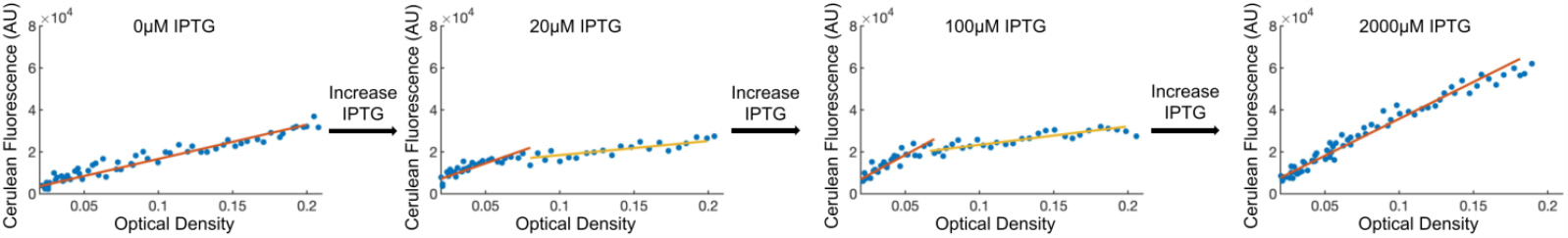
Excision Rate Can Have Bimodal Slope During Exponential Growth. Determination of average excision rate per cell as the slope of mCerulean3 fluorescence versus optical density. For intermediate IPTG concentrations (e.g., 20 μM and 100 μM shown here), the response can appear with bimodal slope. In such cases, the integral of fluorescence versus OD (Cumulative Cerulean) is calculated as given in eq. S3.2.3, with *m*_1_ and *m*_2_ determined as the slopes of the red and yellow lines, respectively.

## SUPPLEMENTAL TABLES

**Table S1.**
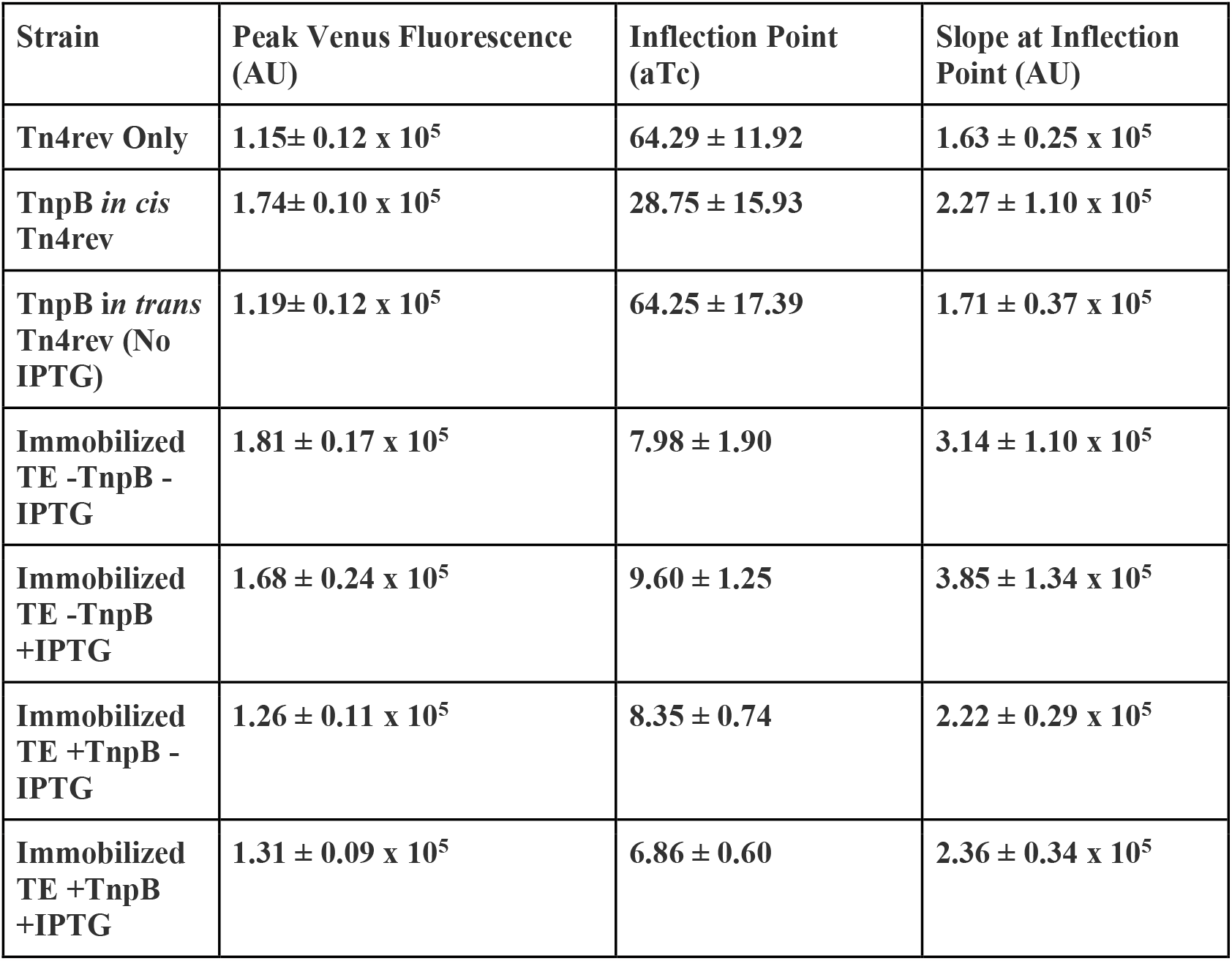
Quantitative Features of Venus-TnpA Induction Response.

**Table S2.**
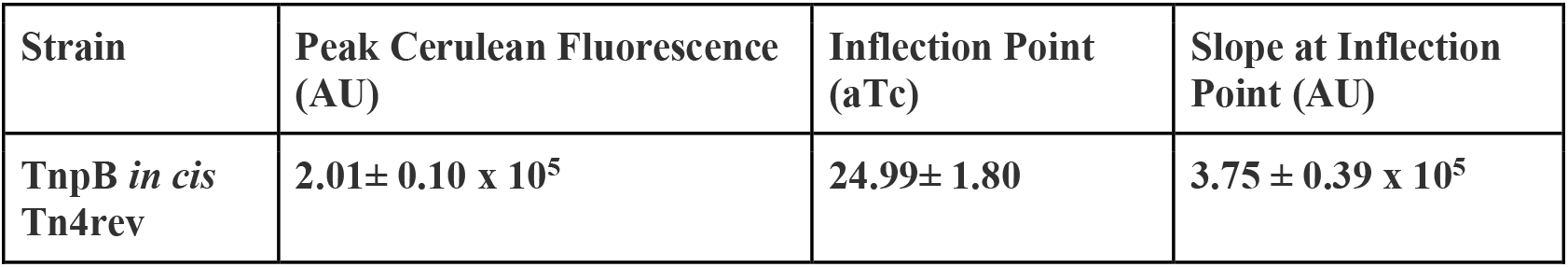
Quantitative Features of Excision Rate Response.

**Table S3:**
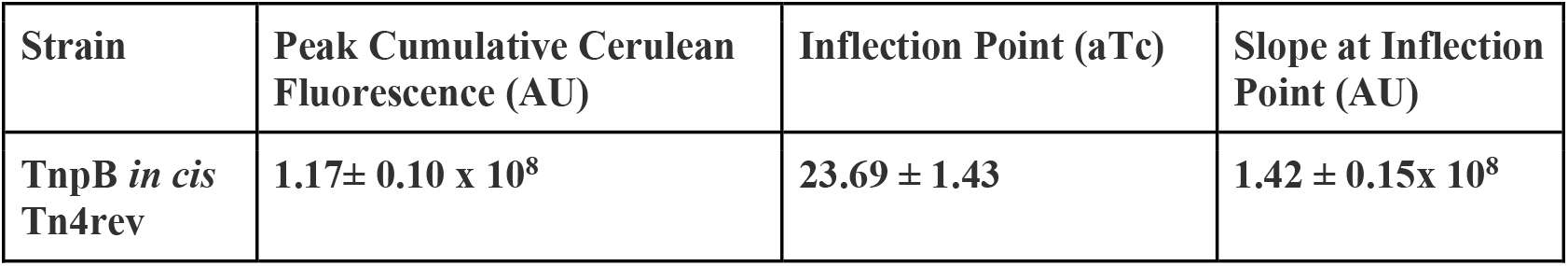
Quantitative Features of Excision Number Response.

**Table S4:**
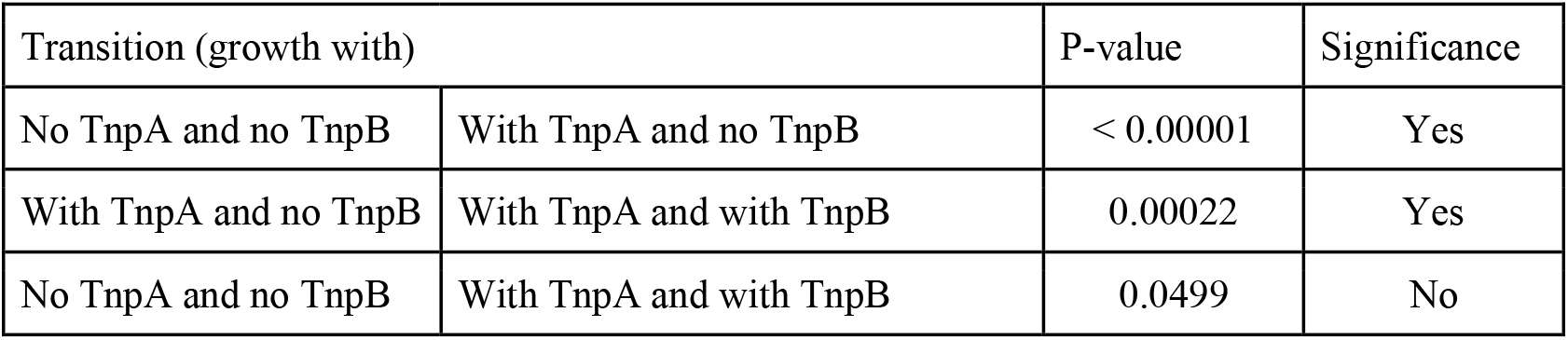
P-values for Changes in Total Transposon Number.

**Table S5:**
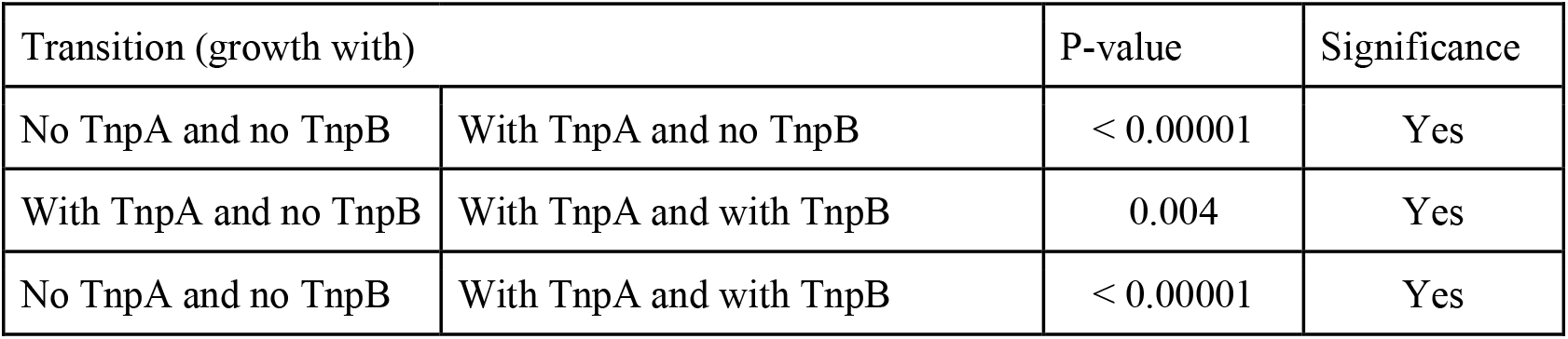
P-values for Changes in Plasmid-Transposon Number.

**Table S6:**
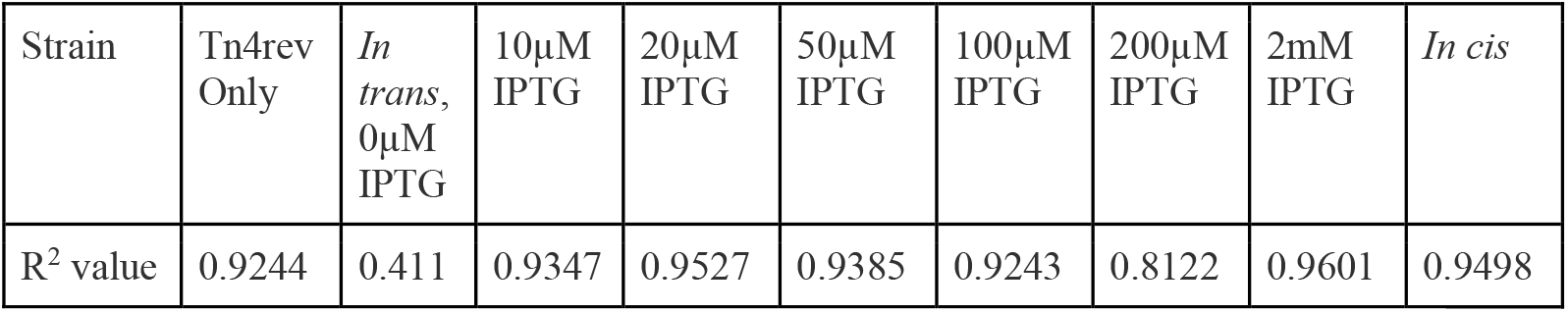
Quality of Fit of Exponential Decay of Growth Rate.

| Strain | Tn4rev Only | *In trans*, 0μM IPTG | 10μM IPTG | 20μM IPTG | 50μM IPTG | 100μM IPTG | 200μM IPTG | 2mM IPTG | *In cis* |
| --- | --- | --- | --- | --- | --- | --- | --- | --- | --- |
| R<sup>2</sup> value | 0.9244 | 0.411 | 0.9347 | 0.9527 | 0.9385 | 0.9243 | 0.8122 | 0.9601 | 0.9498 |

**Table S7:**
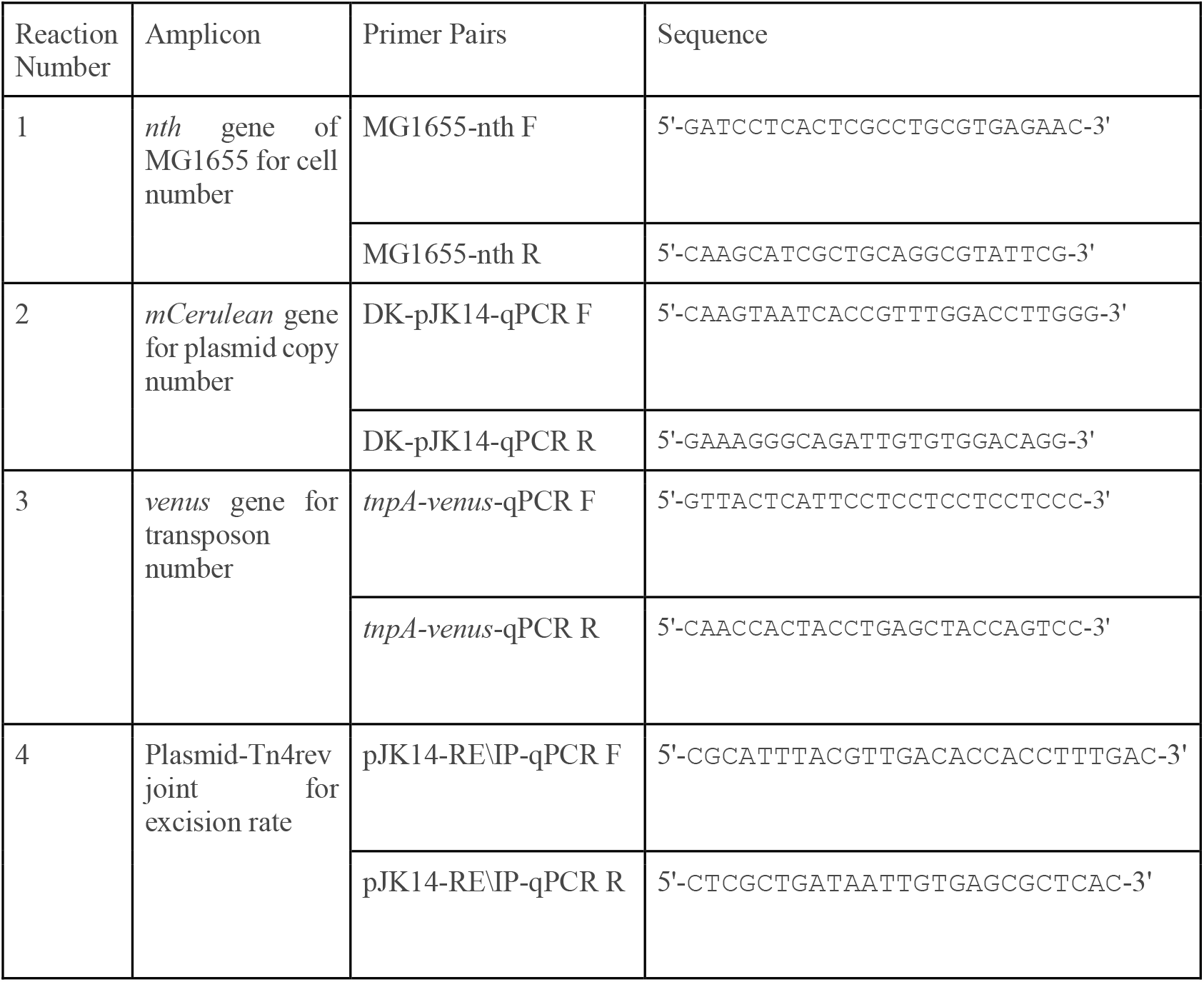
Primer Sequences for qPCR.

## REFERENCES

(1) Shmakov, S.; Smargon, A.; Scott, D.; Cox, D.; Pyzocha, N.; Yan, W.; Abudayyeh, O. O.; Gootenberg, J. S.; Makarova, K. S.; Wolf, Y. I.; Severinov, K.; Zhang, F.; Koonin, E. V. Diversity and Evolution of Class 2 CRISPR–Cas Systems. Nat. Rev. Microbiol. 2017, 15 (3), 169–182. 10.1038/nrmicro.2016.184.

(2) Bao, W.; Jurka, J. Homologues of Bacterial TnpB_IS605 Are Widespread in Diverse Eukaryotic Transposable Elements. Mob. DNA 2013, 4, 12. 10.1186/1759-8753-4-12.

(3) Altae-Tran, H.; Kannan, S.; Demircioglu, F. E.; Oshiro, R.; Nety, S. P.; McKay, L. J.; Dlakić, M.; Inskeep, W. P.; Makarova, K. S.; Macrae, R. K.; Koonin, E. V.; Zhang, F. The Widespread IS200/IS605 Transposon Family Encodes Diverse Programmable RNA-Guided Endonucleases. Science 2021, 374 (6563), 57–65. 10.1126/science.abj6856.

(4) Karvelis, T.; Druteika, G.; Bigelyte, G.; Budre, K.; Zedaveinyte, R.; Silanskas, A.; Kazlauskas, D.; Venclovas, Č.; Siksnys, V. Transposon-Associated TnpB Is a Programmable RNA-Guided DNA Endonuclease. Nature 2021, 599 (7886), 692–696. 10.1038/s41586-021-04058-1.

(5) Pasternak, C.; Dulermo, R.; Ton-Hoang, B.; Debuchy, R.; Siguier, P.; Coste, G.; Chandler, M.; Sommer, S. ISDra2 Transposition in Deinococcus Radiodurans Is Downregulated by TnpB. Mol. Microbiol. 2013, 88 (2), 443–455. 10.1111/mmi.12194.

(6) McClintock, B. The Origin and Behavior of Mutable Loci in Maize. Proc. Natl. Acad. Sci. U. S. A. 1950, 36 (6), 344–355.

(7) Belancio, V. P.; Deininger, P. L.; Roy-Engel, A. M. LINE Dancing in the Human Genome: Transposable Elements and Disease. Genome Med. 2009, 1 (10), 97. 10.1186/gm97.

(8) Callinan, P. A.; Batzer, M. A. Retrotransposable Elements and Human Disease. In Genome Dynamics; Volff, J.-N., Ed.; KARGER: Basel, 2006; pp 104–115. 10.1159/000092503.

(9) Chen, J.-M.; Stenson, P. D.; Cooper, D. N.; Férec, C. A Systematic Analysis of LINE-1 Endonuclease-Dependent Retrotranspositional Events Causing Human Genetic Disease. Hum. Genet. 2005, 117 (5), 411–427. 10.1007/s00439-005-1321-0.

(10) Deininger, P. L.; Batzer, M. A. Alu Repeats and Human Disease. Mol. Genet. Metab. 1999, 67 (3), 183–193. 10.1006/mgme.1999.2864.

(11) Kazazian, H. H.; Wong, C.; Youssoufian, H.; Scott, A. F.; Phillips, D. G.; Antonarakis, S. E. Haemophilia A Resulting from de Novo Insertion of L1 Sequences Represents a Novel Mechanism for Mutation in Man. Nature 1988, 332 (6160), 164–166. 10.1038/332164a0.

(12) O’Donnell, K. A.; Burns, K. H. Mobilizing Diversity: Transposable Element Insertions in Genetic Variation and Disease. Mob. DNA 2010, 1 (1), 21. 10.1186/1759-8753-1-21.

(13) Bodega, B.; Orlando, V. Repetitive Elements Dynamics in Cell Identity Programming, Maintenance and Disease. Curr. Opin. Cell Biol. 2014, 31, 67–73. 10.1016/j.ceb.2014.09.002.

(14) Coufal, N. G.; Garcia-Perez, J. L.; Peng, G. E.; Yeo, G. W.; Mu, Y.; Lovci, M. T.; Morell, M.; O’Shea, K. S.; Moran, J. V.; Gage, F. H. L1 Retrotransposition in Human Neural Progenitor Cells. Nature 2009, 460 (7259), 1127–1131. 10.1038/nature08248.

(15) Anwar, S. L.; Wulaningsih, W.; Lehmann, U. Transposable Elements in Human Cancer: Causes and Consequences of Deregulation. Int. J. Mol. Sci. 2017, 18 (5). 10.3390/ijms18050974.

(16) Burns, K. H. Transposable Elements in Cancer. Nat. Rev. Cancer 2017, 17 (7), 415–424. 10.1038/nrc.2017.35.

(17) Kano, H.; Godoy, I.; Courtney, C.; Vetter, M. R.; Gerton, G. L.; Ostertag, E. M.; Kazazian, H. H. L1 Retrotransposition Occurs Mainly in Embryogenesis and Creates Somatic Mosaicism. Genes Dev. 2009, 23 (11), 1303–1312. 10.1101/gad.1803909.

(18) Schneider, D.; Lenski, R. E. Dynamics of Insertion Sequence Elements during Experimental Evolution of Bacteria. Res. Microbiol. 2004, 155 (5), 319–327. 10.1016/j.resmic.2003.12.008.

(19) Chao, L.; Vargas, C.; Spear, B. B.; Cox, E. C. Transposable Elements as Mutator Genes in Evolution. Nature 1983, 303 (5918), 633–635. 10.1038/303633a0.

(20) Modzelewski, A. J.; Shao, W.; Chen, J.; Lee, A.; Qi, X.; Noon, M.; Tjokro, K.; Sales, G.; Biton, A.; Anand, A.; Speed, T. P.; Xuan, Z.; Wang, T.; Risso, D.; He, L. A Mouse-Specific Retrotransposon Drives a Conserved Cdk2ap1 Isoform Essential for Development. Cell 2021, 184 (22), 5541-5558.e22. 10.1016/j.cell.2021.09.021.

(21) Cordaux, R.; Hedges, D. J.; Herke, S. W.; Batzer, M. A. Estimating the Retrotransposition Rate of Human Alu Elements. Gene 2006, 373, 134–137. 10.1016/j.gene.2006.01.019.

(22) Mills, R. E.; Bennett, E. A.; Iskow, R. C.; Luttig, C. T.; Tsui, C.; Pittard, W. S.; Devine, S. E. Recently Mobilized Transposons in the Human and Chimpanzee Genomes. Am. J. Hum. Genet. 2006, 78 (4), 671–679.

(23) Kim Neil H.; Lee Gloria; Sherer Nicholas A.; Martini K. Michael; Goldenfeld Nigel; Kuhlman Thomas E. Real-Time Transposable Element Activity in Individual Live Cells. Proc. Natl. Acad. Sci. 2016, 113 (26), 7278–7283. 10.1073/pnas.1601833113.

(24) He, S.; Corneloup, A.; Guynet, C.; Lavatine, L.; Caumont-Sarcos, A.; Siguier, P.; Marty, B.; Dyda, F.; Chandler, M.; Ton Hoang, B. The IS200/IS605 Family and “Peel and Paste” Single-Strand Transposition Mechanism. Microbiol. Spectr. 2015, 3 (4). 10.1128/microbiolspec.MDNA3-0039-2014.

(25) Ton-Hoang, B.; Pasternak, C.; Siguier, P.; Guynet, C.; Hickman, A. B.; Dyda, F.; Sommer, S.; Chandler, M. Single-Stranded DNA Transposition Is Coupled to Host Replication. Cell 2010, 142 (3), 398–408. 10.1016/j.cell.2010.06.034.

(26) Guynet, C.; Hickman, A. B.; Barabas, O.; Dyda, F.; Chandler, M.; Ton-Hoang, B. In Vitro Reconstitution of a Single-Stranded Transposition Mechanism of IS608. Mol. Cell 2008, 29 (3), 302–312. 10.1016/j.molcel.2007.12.008.

(27) Ton-Hoang, B.; Guynet, C.; Ronning, D. R.; Cointin-Marty, B.; Dyda, F.; Chandler, M. Transposition of ISHp608, Member of an Unusual Family of Bacterial Insertion Sequences. EMBO J. 2005, 24 (18), 3325–3338. 10.1038/sj.emboj.7600787.

(28) Barabas, O.; Ronning, D. R.; Guynet, C.; Hickman, A. B.; Ton-Hoang, B.; Chandler, M.; Dyda, F. Mechanism of IS200/IS605 Family DNA Transposases: Activation and Transposon-Directed Target Site Selection. Cell 2008, 132 (2), 208–220. 10.1016/j.cell.2007.12.029.

(29) He, S.; Guynet, C.; Siguier, P.; Hickman, A. B.; Dyda, F.; Chandler, M.; Ton-Hoang, B. IS200/IS605 Family Single-Strand Transposition: Mechanism of IS608 Strand Transfer. Nucleic Acids Res. 2013, 41 (5), 3302–3313. 10.1093/nar/gkt014.

(30) Reconstitution of a functional IS608 single-strand transpososome: role of non-canonical base pairing | Nucleic Acids Research | Oxford Academic. https://academic.oup.com/nar/article/39/19/8503/1179505?login=true (accessed 2022-03-07).

(31) Kersulyte, D.; Velapatiño, B.; Dailide, G.; Mukhopadhyay, A. K.; Ito, Y.; Cahuayme, L.; Parkinson, A. J.; Gilman, R. H.; Berg, D. E. Transposable Element ISHp608 of Helicobacter Pylori: Nonrandom Geographic Distribution, Functional Organization, and Insertion Specificity. J. Bacteriol. 2002, 184 (4), 992–1002. 10.1128/jb.184.4.992-1002.2002.

(32) Kersulyte, D.; Mukhopadhyay, A. K.; Shirai, M.; Nakazawa, T.; Berg, D. E. Functional Organization and Insertion Specificity of IS607, a Chimeric Element of Helicobacter Pylori. J. Bacteriol. 2000, 182 (19), 5300–5308. 10.1128/JB.182.19.5300-5308.2000.

(33) Siguier, P.; Gourbeyre, E.; Chandler, M. Bacterial Insertion Sequences: Their Genomic Impact and Diversity. FEMS Microbiol. Rev. 2014, 38 (5), 865–891. 10.1111/1574-6976.12067.

(34) Kapitonov, V. V.; Makarova, K. S.; Koonin, E. V. ISC, a Novel Group of Bacterial and Archaeal DNA Transposons That Encode Cas9 Homologs. J. Bacteriol. 2016, 198 (5), 797–807. 10.1128/JB.00783-15.

(35) Shmakov, S.; Abudayyeh, O. O.; Makarova, K. S.; Wolf, Y. I.; Gootenberg, J. S.; Semenova, E.; Minakhin, L.; Joung, J.; Konermann, S.; Severinov, K.; Zhang, F.; Koonin, E. V. Discovery and Functional Characterization of Diverse Class 2 CRISPR-Cas Systems. Mol. Cell 2015, 60 (3), 385–397. 10.1016/j.molcel.2015.10.008.

(36) Lutz, R.; Bujard, H. Independent and Tight Regulation of Transcriptional Units in Escherichia Coli Via the LacR/O, the TetR/O and AraC/I1-I2 Regulatory Elements. Nucleic Acids Res. 1997, 25 (6), 1203–1210. 10.1093/nar/25.6.1203.

(37) Calos, M. P.; Miller, J. H. The DNA Sequence Change Resulting from the IQ1mutation, Which Greatly Increases Promoter Strength. Mol. Gen. Genet. MGG 1981, 183 (3), 559–560. 10.1007/BF00268783.

(38) Markwardt, M. L.; Kremers, G.-J.; Kraft, C. A.; Ray, K.; Cranfill, P. J. C.; Wilson, K. A.; Day, R. N.; Wachter, R. M.; Davidson, M. W.; Rizzo, M. A. An Improved Cerulean Fluorescent Protein with Enhanced Brightness and Reduced Reversible Photoswitching. PLoS ONE 2011, 6 (3), e17896. 10.1371/journal.pone.0017896.

(39) Real-Time Quantification of the Effects of IS200/IS605 Family-Associated TnpB on Transposon Activity. https://www.jove.com/t/64825/real-time-quantification-of-the-effects-of-is200-is605-family-associated-tnpb-on-transposon-activity (accessed 2023-01-26).

(40) Weinreich, M. D.; Gasch, A.; Reznikoff, W. S. Evidence That the Cis Preference of the Tn5 Transposase Is Caused by Nonproductive Multimerization. Genes Dev. 1994, 8 (19), 2363–2374. 10.1101/gad.8.19.2363.

(41) Cerveau, N.; Leclercq, S.; Leroy, E.; Bouchon, D.; Cordaux, R. Short- and Long-Term Evolutionary Dynamics of Bacterial Insertion Sequences: Insights from Wolbachia Endosymbionts. Genome Biol. Evol. 2011, 3, 1175–1186. 10.1093/gbe/evr096.

(42) Wagner, A. Periodic Extinctions of Transposable Elements in Bacterial Lineages: Evidence from Intragenomic Variation in Multiple Genomes. Mol. Biol. Evol. 2006, 23 (4), 723–733. 10.1093/molbev/msj085.

(43) Koonin, E. V.; Makarova, K. S. Mobile Genetic Elements and Evolution of CRISPR-Cas Systems: All the Way There and Back. Genome Biol. Evol. 2017, 9 (10), 2812–2825. 10.1093/gbe/evx192.

(44) Krupovic, M.; Makarova, K. S.; Forterre, P.; Prangishvili, D.; Koonin, E. V. Casposons: A New Superfamily of Self-Synthesizing DNA Transposons at the Origin of Prokaryotic CRISPR-Cas Immunity. BMC Biol. 2014, 12 (1), 36. 10.1186/1741-7007-12-36.

(45) Koonin, E. V.; Krupovic, M. Evolution of Adaptive Immunity from Transposable Elements Combined with Innate Immune Systems. Nat. Rev. Genet. 2015, 16 (3), 184–192. 10.1038/nrg3859.

(46) Gene Location and DNA Density Determine Transcription Factor Distributions in Escherichia Coli. Mol. Syst. Biol. 2012, 8 (1), 610. 10.1038/msb.2012.42.

(47) Kuhlman, T. E.; Cox, E. C. DNA-Binding-Protein Inhomogeneity in $E.$ Coli Modeled as Biphasic Facilitated Diffusion. Phys. Rev. E 2013, 88 (2), 022701. 10.1103/PhysRevE.88.022701.

(48) Kuhlman, T. E.; Cox, E. C. Site-Specific Chromosomal Integration of Large Synthetic Constructs. Nucleic Acids Res. 2010, 38 (6), e92. 10.1093/nar/gkp1193.

(49) Kuhlman, T. E.; Cox, E. C. A Place for Everything. Bioeng. Bugs 2010, 1 (4), 296–299. 10.4161/bbug.1.4.12386.

(50) Tas, H.; Nguyen, C. T.; Patel, R.; Kim, N. H.; Kuhlman, T. E. An Integrated System for Precise Genome Modification in Escherichia Coli. PloS One 2015, 10 (9), e0136963. 10.1371/journal.pone.0136963.

(51) Kinney, J. B.; Murugan, A.; Callan, C. G.; Cox, E. C. Using Deep Sequencing to Characterize the Biophysical Mechanism of a Transcriptional Regulatory Sequence. Proc. Natl. Acad. Sci. 2010, 107 (20), 9158–9163. 10.1073/pnas.1004290107.

(52) del Solar, G.; Giraldo, R.; Ruiz-Echevarría, M. J.; Espinosa, M.; Díaz-Orejas, R. Replication and Control of Circular Bacterial Plasmids. Microbiol. Mol. Biol. Rev. MMBR 1998, 62 (2), 434–464. 10.1128/MMBR.62.2.434-464.1998.

(53) Tucker, W. T.; Miller, C. A.; Cohen, S. N. Structural and Functional Analysis of the Par Region of the PSC 10 1 Plasmid. Cell 1984, 38 (1), 191–201. 10.1016/0092-8674(84)90540-3.

## References

(1) Kim Neil H.;Lee Gloria; Sherer Nicholas A.; Martini K. Michael; Goldenfeld Nigel; Kuhlman Thomas E. Real-Time Transposable Element Activity in Individual Live Cells. Proc. Natl. Acad. Sci. 2016, 113 (26), 7278–7283. 10.1073/pnas.1601833113.

(2) Gene Location and DNA Density Determine Transcription Factor Distributions in Escherichia Coli. Mol. Syst. Biol. 2012, 8 (1), 610. 10.1038/msb.2012.42.

(3) Kuhlman, T. E.; Cox, E. C. Site-Specific Chromosomal Integration of Large Synthetic Constructs. Nucleic Acids Res. 2010, 38 (6), e92. 10.1093/nar/gkp1193.

(4) Tas, H.; Nguyen, C. T.; Patel, R.; Kim, N. H.; Kuhlman, T. E. An Integrated System for Precise Genome Modification in Escherichia Coli. PloS One 2015, 10 (9), e0136963. 10.1371/journal.pone.0136963.

