## Supplementary Information for "IS200/IS605 Family-Associated TnpB Increases Transposon Activity and Retention"

#### SI. SUPPLEMENTARY ANALYSIS

##### SI.i. Introduction of TnpB to the Inducible Transposon

TnpB is translationally fused to mCherry and introduced to the transposon *in trans* and individually induced with IPTG (0 µM, 10 µM, 20 µM, 50 µM, 100 µM, 200 µM & 2000 µM). Levels of mCherry-TnpB as measured by mCherry fluorescence increase upon titration with IPTG (**Supplementary Fig. S1B**). The data are fit to a Hill function of the form

$$H\left(\left[IPTG\right]\right)=E_{\max}\left(\frac{\left(\left[IPTG\right]/K_{IPTG}\right)^{n_{IPTG}}}{1+\left(\left[IPTG\right]/K_{IPTG}\right)^{n_{IPTG}}}\right)+E_{\min}, \qquad \text{(S1.1.1)}$$

as described in the Methods section **Quantification of Fluorescence Per Cell**. Here, $E_{max}$ is the overall scaling factor, $E_{min}$ is the minimum value of the fit, $n_{IPTG}$ is the Hill coefficient and $K_{IPTG}$ is the LacI-IPTG dissociation constant. We find that $K_{IPTG}$ = 61.53 µM and $n_{IPTG}$ = 1.03.

Similarly, when TnpB is translationally fused to mCherry and introduced to the transposon *in cis* it is induced with aTc (0 ng/mL, 1 ng/mL, 2 ng/mL, 3 ng/mL, 5 ng/mL, 9 ng/mL, 20 ng/mL, 30 ng/mL, 50 ng/mL, 80 ng/mL, 100 ng/mL and 200 ng/mL). Levels of mCherry-TnpB as measured by mCherry fluorescence increase upon titration with aTc (**Supplementary Fig. S1C**). The data are fit to a Hill function of the form

$$H\left(\left[aTc\right]\right)=E_{\max}\left(\frac{\left(\left[aTc\right]/K_{aTc}\right)^{n_{aTc}}}{1+\left(\left[aTc\right]/K_{aTc}\right)^{n_{aTc}}}\right)+E_{\min}, \qquad \text{(S1.1.2)}$$

where parameters are defined similar to those in eq. (S1.1.1). We find that $K_{aTc}$ = 285.77 ng/mL and $n_{aTc}$ = 0.98.

$$\log\left(\frac{F(t)}{OD(t)}\right) = -\beta\left[\log\left(OD(t=0)\right) + gt\right] + A, \tag{S1.6.1}$$

where $g$ is the growth rate of the cells (determined as described in Methods), $t$ is time in seconds, and $A$ is a constant. Here, the fluorescence per optical density, $\mathcal{F}_C$(t), decays as a function of time as follows:

$$\mathcal{F}_C(t) = \mathcal{F}_C(t=0)\left[OD(t=0)\right]^{-\beta} e^{-\beta g t}. \tag{S1.6.2}$$

These exponent values, $\beta g$, are then normalized to values for the negative control strain and plotted (**Supplementary Fig. S6**). For lower concentrations of both proteins, the number of transposase molecules decays at a higher rate over time. Increase in TnpA and TnpB both result in maintenance of transposase concentration due to a lower decay rate over time. Further, TnpA and TnpB have a compounding effect on maintenance of TEs.

$$q = \binom{L}{0} w^0 (1-w)^L = \exp\left[-\ln\left(\frac{1}{1-w}\right)L\right]. \tag{S1.8.1}$$

In our growth experiments, an exponentially growing individual cell, in the absence of integrations, will produce $g_0 dt$ new individuals, in a time interval $dt$. This leads to a simple model of exponential growth of the form $\frac{dx}{dt} = g_0 x$. If we consider a binary model with a population $x$ of normal cells and a population $y$ of cells with no growth due to integrations, an individual of $x$ will still produce $g_0 dt$ new individuals but only a fraction $q$ of these will be able to grow. This leads to the population model:

$$\frac{dx}{dt} = qg_0 x, \quad \frac{dy}{dt} = (1-q)g_0 x. \tag{S1.8.2}$$

The total population of cells in this model grows as $x_0 + y_0 + \frac{x_0}{q}\left[\exp(qg_0 t) - 1\right]$. Thus, the measured growth rate would be $qg_0$ and the normalized growth rate is just $q$. We fit eq. (S1.8.1) to the form $\exp[-bL]$ and make the identification $b = -\ln[1-w]$, which means $b \approx w$ for $w \ll 1$. That is, $b$ is approximately equal to the probability of a transposon transcript integrating and disrupting growth. Moreover, we expect that the rate of obtaining integrations affecting growth, $w$, is proportional to the overall rate of integration, $\mu$. Consequently, this simple binary model recapitulates the exponential dependence of the growth rate on the number of transcripts and demonstrates that the exponential dependence implies that the growth defect, $b$, is expected to be directly coupled to the integration rate, $\mu$.

More complex models of the impact of transposable element integration can be developed, with more than two sub-populations and more nuanced assumptions about the physiological effects. But we find that the dynamics of these models reduce to that of the two-rate model presented above, with renormalized parameters. An example of one such model is as follows. Let the number of cells with no chromosomal integrations harming their growth be $N_0$, the population of cells with one integrant be $N_1$, and so forth. Then a set of differential equations describing the population dynamics in exponential growth with growth rate $g_0$ is

$$\frac{dN_x}{dt} = g_0 f(x)(1-\mu)N_x + g_0 f(x-1)\mu N_{x-1}, \tag{S1.8.3}$$

where $f(x)$ is a monotonically decreasing function describing the inhibition of cell growth due to gene disruption by integrations, $\mu$ is the mutation rate, and the index $x$ runs from 0 to some integer $x_{max}$ such that the number of integrants is so high the cell cannot function and dies. Making the substitution $(1-\mu)=q$, we have

$$\mu' = \begin{cases} \mu \frac{T}{X_0}(1 + [TnpB]) & for\, \mu \cdot T \leq X_0 \\ 1 + [TnpB] & for\, \mu \cdot T > X_0 \end{cases}, \tag{S2.1.2}$$

where [TnpB] is a measure of the concentration of TnpB, 0 ≤ [TnpB] ≤ 1 and [TnpB] = 1 corresponds to saturation of the cells with TnpB.

During each cycle, each transposon has a probability $\chi$' of excising per generation, leaving behind one empty plasmid strand, and one with the DNA complementary to the TE. These excised transposons then have a probability $\mu$' of successfully reinserting at a new location per generation. The schematic in **Fig.4A** provides an illustration of how the cell will evolve over time. The following coupled differential equations, along with the expressions above for $\chi$' and $\mu$', are concurrently solved numerically to track the changes in the transposition rates and the plasmid and transposon numbers over time t.

Number of plasmids with transposons *X*(*t*):

$$\frac{dX(t)}{dt} = -\frac{\chi'}{2}X(t) + C_{Hom}[TnpB]\left(X_0 - X(t) + \frac{\chi'}{2}X(t)\right) \quad \text{(S2.1.3)}$$

Number of reinserted transposons *Ch*(t):

$$\frac{dCh(t)}{dt} = \frac{\mu'\chi'}{2}X(t) - \frac{(1-\mu')\chi'}{2}Ch(t) + \frac{C_{hom}[TnpB]\chi'}{2}Ch(t) \quad \text{(S2.1.4)}$$

Number of plasmids without transposons *Y*(t):

$$\frac{dY(t)}{dt} = -\frac{dX(t)}{dt} \quad \text{(S2.1.5)}$$

Total number of transposons, *T*(t):

$$\frac{dT(t)}{dt} = \frac{dX(t)}{dt} + \frac{dCh(t)}{dt} \quad \text{(S2.1.6)}$$

or

$$\frac{dT(t)}{dt} = \frac{(1-\mu')\chi'}{2}(X(t) + Ch(t)) + C_{hom}[TnpB]\left(X_0 - X(t) + \frac{\chi'}{2}(X(t) + Ch(t))\right) \quad \text{(S2.1.7)}$$

$$H\left(\left[inducer\right]\right)=E_{\max}\left(\frac{\left(\left[inducer\right]/K_{inducer}\right)^{n_{inducer}}}{1+\left(\left[inducer\right]/K_{inducer}\right)^{n_{inducer}}}\right)+E_{\min}, \tag{S3.1.1}$$

where $E_{max}$ is the overall scaling factor, $E_{min}$ is the minimum value of the fit, $n_{inducer}$ is the Hill coefficient and $K_{inducer}$ is the activation coefficient. The quantitative features of the responses are extracted. We use the inflection point to compare relative initiation points of the Hill fits. The inflection point is calculated using the following equation

$$x^{*}=\left(\frac{n_{inducer}-1}{n_{inducer}+1}\right)^{\frac{1}{n_{inducer}}}K_{inducer}. \tag{S3.1.2}$$

The inflection point corresponds to the following value of the Hill function,

$$H\left(x^{*}\right)=\frac{E_{\max}}{2}\left(1-\frac{1}{n_{inducer}}\right). \tag{S3.1.3}$$

The slope at the inflection point is used as a measure of the sensitivity of the system to induction and is calculated using the equation

$$S\left(x^{*}\right)=\left.\frac{dH}{dx}\right|_{x=x^{*}}=\left(\frac{n_{inducer}^{2}-1}{2n_{inducer}}\right)\left(\frac{n_{inducer}-1}{n_{inducer}+1}\right)^{\frac{1}{n_{inducer}}}E_{\max}. \tag{S3.1.4}$$

The error in the location of the inflection is determined using the 95% confidence interval of the Hill fit variables and the following equation

$$\delta x^{*}=x^{*}\sqrt{\left(\frac{2}{n_{inducer}\left(n_{inducer}^{2}-1\right)}-\frac{\log\left(\frac{n_{inducer}-1}{n_{inducer}+1}\right)}{n_{inducer}^{2}}\right)^{2}\delta n_{inducer}^{2}+\frac{\delta K_{inducer}^{2}}{K_{inducer}^{2}}}. \tag{S3.1.5}$$

where $\delta n_{inducer}$ is the standard error of the mean of the Hill coefficient. The error in the slope of the inflection is determined using the 95% confidence interval of the Hill fit variables and the following equation

$$\delta S\left(x^{*}\right)=S^{*}\sqrt{\left(\frac{1}{n_{inducer}}+\frac{1}{n_{inducer}^{2}}\log\left(\frac{n_{inducer}-1}{n_{inducer}+1}\right)\right)^{2}\delta n_{inducer}^{2}+\frac{\delta E_{\max}^{2}}{E_{\max}^{2}}}. \tag{S3.1.6}$$

**SIII.ii. Calculation of Cumulative Cerulean Fluorescence Per Cell**

In previous work, we demonstrate that excision events are followed by spikes of Cerulean fluorescence. Thus, we measure excision rate per cell as the slope of Cerulean fluorescence vs. optical density, as described in the section **Quantification of Fluorescence per Cell**. In **Supplementary Fig. S7**, we illustrate the bimodal nature of the slope of excision rate that we observe during the exponential growth phase of *E. coli*. In **Supplementary Fig. S7**, we plot the initial excision rate, $m_1$, starting at the lowest OD~0.01 until the slope changes (green selection region of schematic) at point OD*. However, to determine the total number of excision events from the plasmids in the exponential phase, we integrate Cerulean fluorescence per cell (AU) over time. We measure $m_2$ to make the following calculation of cumulative Cerulean or total plasmid excision events during exponential growth.

Cerulean Fluorescence per cell is the rate of excision from plasmids,

$$\frac{Cerulean}{cell}=r_c\frac{d\left(Excised\ Plasmids\right)}{dt}, \tag{S3.2.1}$$

where, $r_c$ is the fluorescence per plasmid-excision event. The integral of Cerulean Fluorescence per cell over time is the total number of excision events from plasmids:

$$\frac{Cumulative\ Cerulean}{cell} = \int \frac{Cerulean}{cell} dt = r_c \left( Excised\ \ Plasmids \right). \tag{S3.2.2}$$

Executing the integral over OD, we obtain

$$\frac{Cumulative\ Cerulean}{cell} = \frac{m_1 - m_2}{g'} \ln\left(OD^*\right) + \frac{m_2}{g'} \ln\left(0.2\right) - \frac{m_1}{g'} \ln\left(0.01\right), \tag{S3.2.3}$$

where $g' = g\ln\left(2\right)$ and $g$ is the growth rate of the cells.

#### SIII.iii. Quantitative-PCR Analysis

Quantitative PCR reactions with all four primer pairs from **Supplementary Table S7** were performed concurrently on the dilution series of cells with reactions performed in quadruplicate. The quantity of cells is conserved between reactions for each dilution factor $d_i$ ($d_i \in [5^1, 5^6]$) and the amplification from the MG1655 *nth* chromosomal locus located near the replication terminus[2–4] allows for a measurement proportional to the number of cells. This allows the following calculation of the relative number of each gene per cell:

$$N\left(n_N\right) = N_0\left(1+\varepsilon_N\right)^{n_N} \tag{S3.3.1}$$

$$K\left(n_K\right) = K_0\left(1+\varepsilon_K\right)^{n_K}\ , \tag{S3.3.2}$$

where $N$ is the number of amplicons in the reaction, $\epsilon_N$ is the efficiency of the reaction, and $n_N$ is the cycle number for qPCR reactions 2-4 (**Supplementary Table S7**): plasmid copy number, total transposon number, or unexcised plasmid number. Here, $K$ is the number of MG1655 *nth* gene amplicons in the reaction (primer pair 1), $\epsilon_K$ is the efficiency of the reaction and $n_K$ is the cycle number. When we consider the ratio of these equations at the threshold cycle number,

$$\frac{N(C_{Ni})}{K(C_{Ki})} = \frac{N_0 (1+\varepsilon_N)^{C_{Ni}}}{K_0 (1+\varepsilon_K)^{C_{Ki}}}. \tag{S3.3.3}$$

Using the same arbitrary threshold line for all reactions to determine the threshold cycle numbers of the *i*th sample ($C_{Ni}/C_{Ki}$, $i$ = 1,2...6) makes the left-hand side of eq. (S3.3.3) equal to one and this gives us a relative value of the initial concentration of the sequence of interest per cell, i.e., $N(C_{Ni}) = K(C_{Ki})$, to obtain

$$\frac{N_0}{K_0} = \frac{(1+\varepsilon_K)^{C_{Ki}}}{(1+\varepsilon_N)^{C_{Ni}}}. \tag{S3.3.4}$$

These relative values are determined for each combination of aTc (0 ng/mL and 200 ng/mL) and IPTG (0 mM and 2 mM) concentration for comparison.

#### SIII.iv. Error Calculation for Quantitative-PCR

Quantitative PCR reactions with all four primer pairs from **Supplementary Table S7** were performed concurrently on the dilution series of cells with reactions performed in quadruplicate. The average relative number of each amplicon per cell for each dilution level, $i$ = 1, 2, ..., 6, was determined from four replicate reactions:

$$\chi_{i,p} = \left(\frac{N_0}{K_0}\right)_{i,p} = \frac{(1+\varepsilon_K)^{\bar{C}_{Ki}}}{(1+\varepsilon_N)^{\bar{C}_{Ni}}}, \tag{S3.4.1}$$

where $p$ =1, …, $P$, is the experiment number, and $\bar{C}_{Ni}$ and $\bar{C}_{Ki}$ are the threshold cycle numbers of the $i^{th}$ dilution level averaged over the four replicates. The standard deviation of the relative number of each amplicon per cell for each dilution level, $i$ = 1, 2, ..., 6, was determined as follows:

$$\sigma_{\chi_{i,p}} = \chi_{i,p}\sqrt{\ln^2(1+\varepsilon_N)\,\sigma^2_{\bar{C}_{Ni}} + \ln^2(1+\varepsilon_K)\,\sigma^2_{\bar{C}_{Ki}}} \tag{S3.4.2}$$

where $\sigma_{\bar{C}_{Ni}}$ and $\sigma_{\bar{C}_{Ki}}$ are the standard deviations of threshold cycle numbers of the four replicates of the $i^{th}$ dilution level. The [$\chi_{i,p}$,$\sigma_{\chi i,p}$ ] values from all experiments for each combination of aTc and IPTG inducer concentrations were then pooled. These pooled values were averaged over all experiments and dilution levels to determine the average amplicon number per cell for cells induced with a particular combination of IPTG and aTc concentration,

$$\bar{\chi}_{IPTG,aTc} = Mean\left(\chi_{i,p}\right) + Mean\left(\sigma_{\chi_{i,p}}\right). \tag{S3.4.3}$$

The standard deviation of the pooled $\chi_{i,p}$ values was determined to be $\sigma_{\chi}$. The total standard deviation of the amplicon number per cell for each IPTG and aTc concentration was calculated by accounting for the propagating error from $\sigma_{\chi i,p}$, the values of which have a standard deviation of $\sigma_{\delta\chi i,p}$. The total standard deviation of each $\bar{\chi}_{IPTG,aTc}$ value is then

$$\sigma_{\chi_{IPTG,aTc}} = \sqrt{\sigma_{\chi}^2 + \sigma_{\delta\chi_{i,p}}^2}. \tag{S3.4.4}$$

The uncertainties were determined by calculating the standard error of the mean:

$$SEM_{\chi_{IPTG,aTc}} = \frac{\sigma_{\chi_{IPTG,aTc}}}{\sqrt{\mathrm{Rep}}}, \tag{S3.4.5}$$

where Rep is the number of successful replicates.

### SUPPLEMENTAL FIGURES

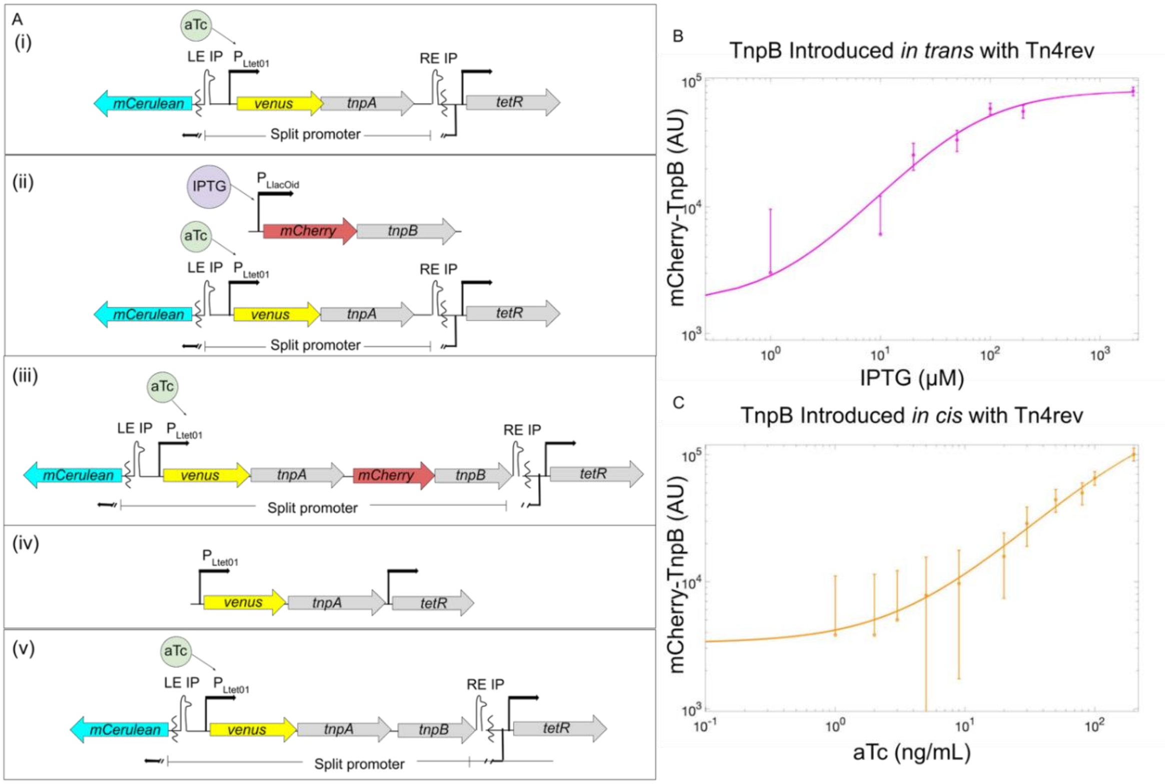


**Figure S1: (A) Genetic Constructs for Inducible and Trackable Transposons:** All transposons constructed for experiments. The strain which has Tn4rev only (i), MG1655 Δ*lac* pJK14-$P_{Ltet-01}$-Tn4rev & pZA31- $P_{LlacOid}$-SmR. TnpB introduced to the transposon *in trans* (ii), MG1655 Δ*lac* pJK14-$P_{Ltet-01}$-*Tn4rev* & pZA31-$P_{LlacOid}$-*mCherry-tnpB* and *in cis* (iii), MG1655 Δ*lac* pJK14-$P_{Ltet-01}$-*tn4rev-mCherry-tnpB* & pZA31- $P_{LlacOid}$-SmR. A version of the transposon (iv) without the excision sites, rendering it immobile. To the immobile transposon, we introduce, *in trans*, both TnpB: CZ071 pJK14-$P_{Ltet-01}$-*tnpA* & pZA31-$P_{LlacOid}$-*mCherry-tnpB*, and its negative control plasmid: CZ071 pJK14-$P_{Ltet-01}$-*tnpA* & pZA31-$P_{LlacOid}$-SmR. A control for the mCherry fusion to TnpB is also constructed (v), MG1655 Δ*lac* pJK14-$P_{Ltet-01}$-*tn4rev-tnpB* & pZA31- $P_{LlacOid}$-SmR. **(B)** TnpB is translationally fused to mCherry, introduced to the transposon *in trans* and individually induced with IPTG (0 μM, 10 μM, 20 μM, 50 μM, 100 μM, 200 μM & 2000 μM). **(C)** TnpB is translationally fused to mCherry and introduced to the transposon *in cis*. mCherry-TnpB and Venus-TnpA are induced simultaneously with aTc (0 ng/mL, 1 ng/mL, 2 ng/mL, 3 ng/mL, 5 ng/mL, 9 ng/mL, 20 ng/mL, 30 ng/mL, 50 ng/mL, 80 ng/mL, 100 ng/mL and 200 ng/mL).

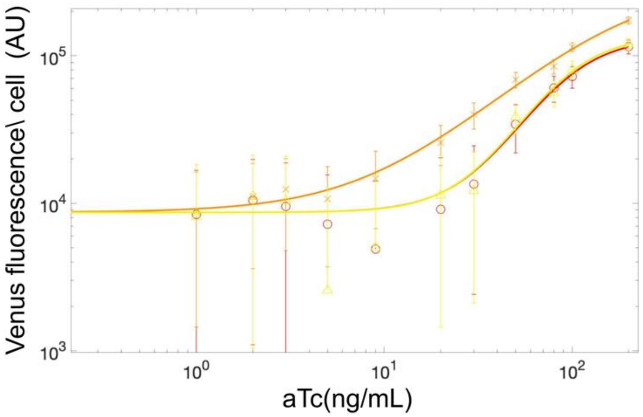


**Figure S2: Quantification of Venus-Transposase Concentration to aTc Induction:** Venus fluorescence data as a function of aTc concentration (0ng/mL, 1ng/mL, 2ng/mL, 3ng/mL, 5ng/mL, 9ng/mL, 20ng/mL, 30ng/mL, 50ng/mL, 80ng/mL, 100ng/mL and 200ng/mL) for **(A)** (i) Tn4rev only strain: MG1655 Δ*lac* pJK14-$P_{Ltet\text{-}01}$-Tn4rev & pZA31-$P_{LlacOid}$-SmR (red, O), (ii) TnpB introduced *in trans* with Tn4rev at [IPTG] = 0: MG1655 Δ*lac* pJK14-$P_{Ltet\text{-}01}$-Tn4rev & pZA31-$P_{LlacOid}$-*mCherry-tnpB* (light green, Δ), (iii) TnpB introduced *in cis* with Tn4rev: MG1655 Δ*lac* pJK14-$P_{Ltet\text{-}01}$-Tn4rev-*mCherry-tnpB* & pZA31-$P_{LlacOid}$-SmR (orange, X).

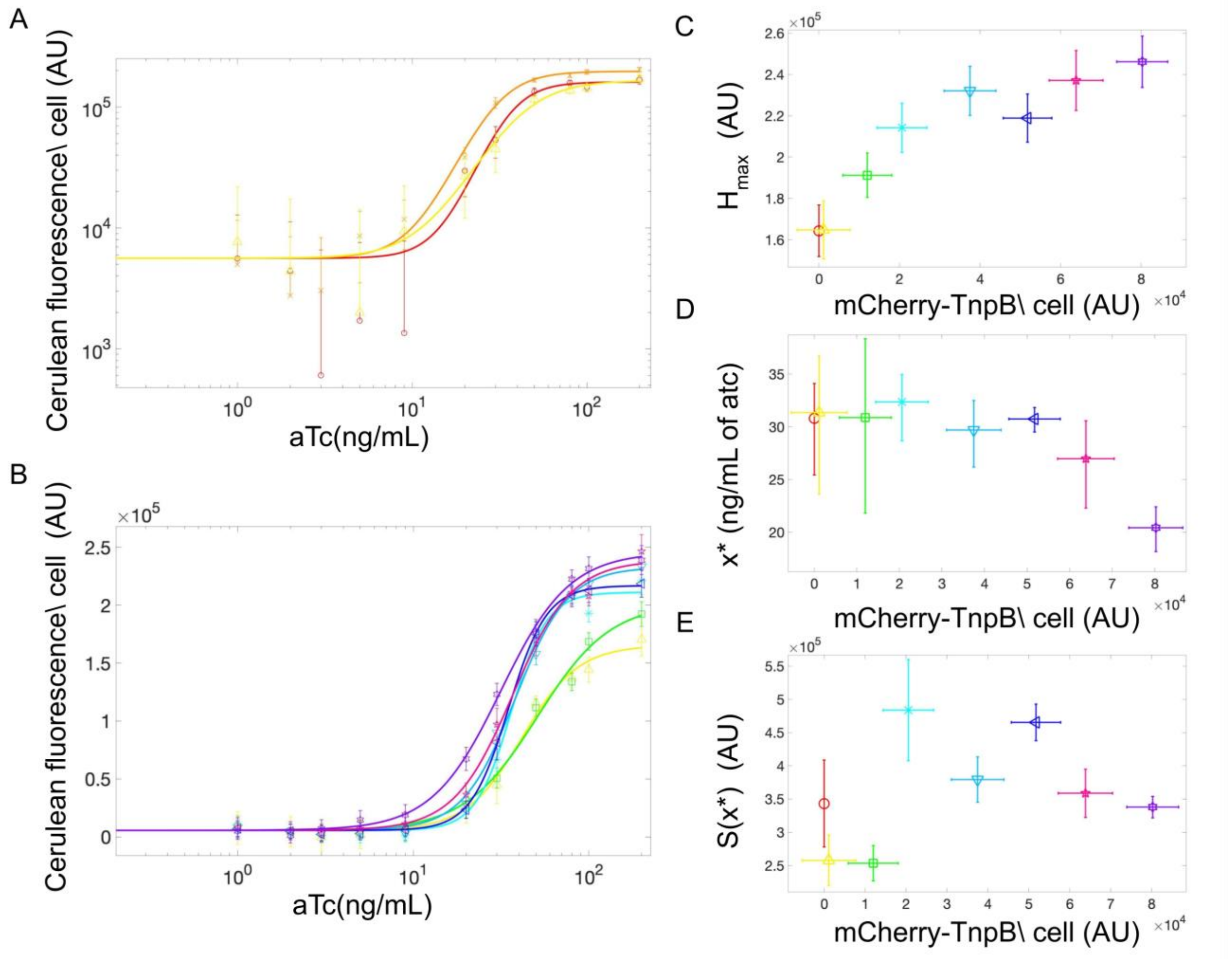


**Figure S3: Titration of Excision Rate with Anhydrotetracycline (aTc): (A,B)** Cerulean fluorescence data for [aTc] = {0ng/mL, 1ng/mL, 2ng/mL, 3ng/mL, 5ng/mL, 9ng/mL, 20ng/mL, 30ng/mL, 50ng/mL, 80ng/mL, 100ng/mL and 200ng/mL} for **(A)** (i) Tn4rev only strain: MG1655 Δ*lac* pJK14-$P_{Ltet\text{-}01}$-Tn4rev & pZA31-$P_{LlacOid}$-SmR (red, ○), (ii) TnpB introduced *in trans* with Tn4rev, [IPTG] = 0: MG1655 Δ*lac* pJK14-$P_{Ltet\text{-}01}$-Tn4rev & pZA31-$P_{LlacOid}$-*mCherry-tnpB* (light green, Δ), (iii) TnpB introduced *in cis* with Tn4rev: MG1655 Δ*lac* pJK14-$P_{Ltet\text{-}01}$-01-Tn4rev-*mCherry-tnpB* & pZA31-$P_{LlacOid}$-SmR (orange, X), and **(B)** TnpB is introduced *in trans* with Tn4rev, MG1655 Δ*lac* pJK14-$P_{Ltet\text{-}01}$-Tn4rev pZA31-$P_{LlacOid}$-*mCherry-tnpB* for different concentrations of IPTG [0µM (yellow, Δ), 10µM (green, □), 20µM (turquoise, ✳), 50µM (teal, ▽), 100µM (navy, ◁), 200µM (magenta, ✋) & 2000µM (violet, ✵)]. Cerulean fluorescence data sets are each fit to a Hill function (same color as data points for each) of the form of eq. **(S1.1.2)** and quantitative features of the responses are extracted and included in **C-E** and **Supplementary Table S2** for the strain with TnpB *in cis* with Tn4rev as it does not have a singular mCherry-TnpB value. **(C)** Higher Cerulean levels with TnpB induction when TnpA is maximally expressed ($H_{max}$). **(D)** Earlier initiation of Cerulean expression (inflection point, $x^*$) for lower aTc concentrations with increased TnpB induction. **(E)** Indiscernible pattern of excision sensitivity to aTc concentration (slope at inflection point $S(x^*)$) with increased TnpB concentration.

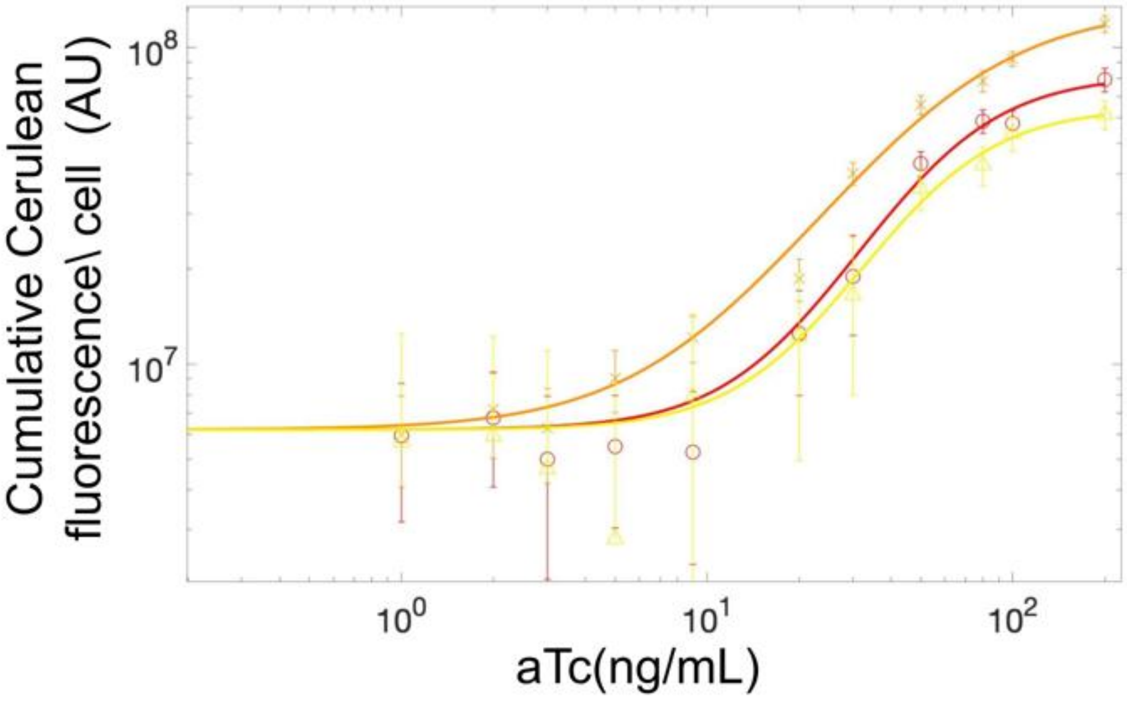


**Figure S4: Titration of Excision Event Number with Anhydrotetracycline (aTc):** Cumulative Cerulean fluorescence data as a function of inducer concentration for (0ng/mL, 1ng/mL, 2ng/mL, 3ng/mL, 5ng/mL, 9ng/mL, 20ng/mL, 30ng/mL, 50ng/mL, 80ng/mL, 100ng/mL and 200ng/mL) for (i) Tn4rev only strain: MG1655 Δ*lac* pJK14-$P_{\textit{Ltet-01}}$-Tn4rev & pZA31-$P_{\textit{LlacOid}}$-SmR (red, O), (ii) TnpB introduced *in trans* with Tn4rev: MG1655 Δ*lac* pJK14-$P_{\textit{Ltet-01}}$-Tn4rev & pZA31-$P_{\textit{LlacOid}}$-*mCherry-tnpB* (light green, Δ), and (iii) TnpB introduced *in cis* with Tn4rev: MG1655 Δ*lac* pJK14-$P_{\textit{Ltet-01}}$-01-Tn4rev-*mCherry-tnpB* & pZA31-$P_{\textit{LlacOid}}$-SmR (orange, X).

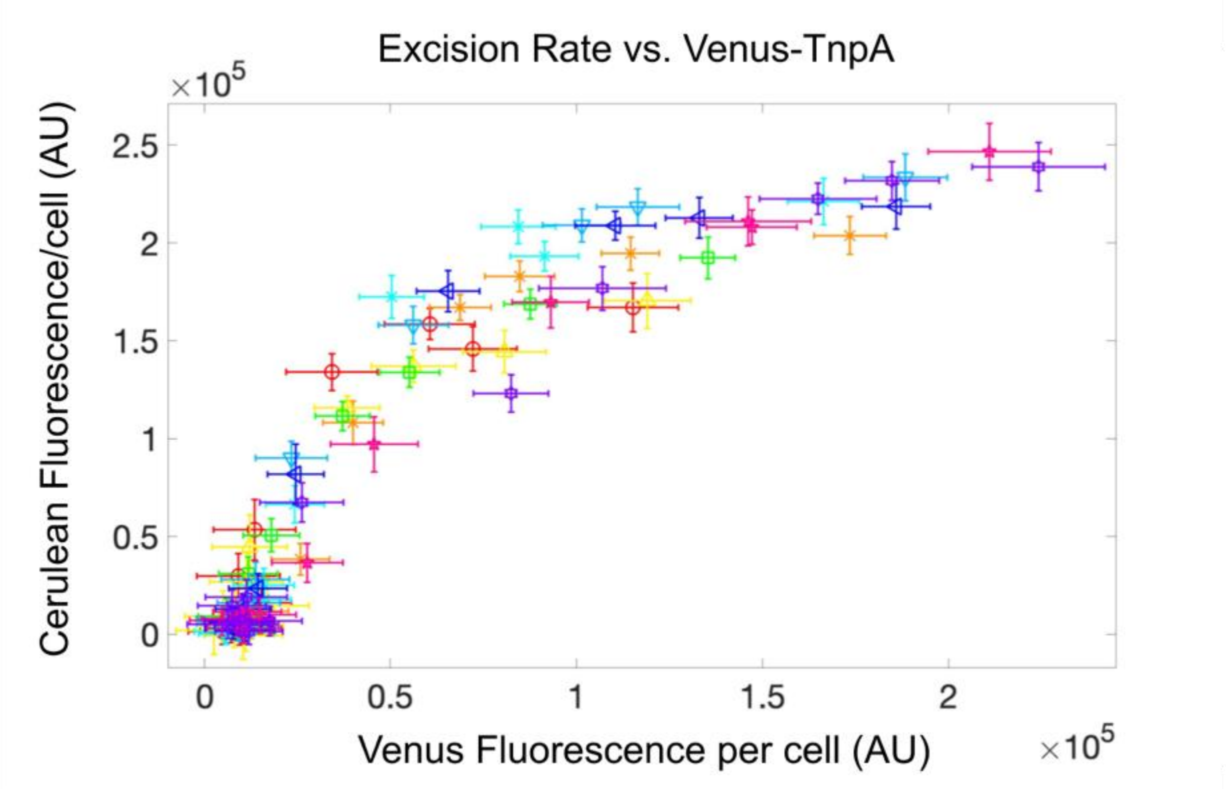


**Figure S5: Excision Rate Increases and Peaks with Transposase Concentration:** Excision rate (Cerulean fluorescence per cell) vs. transposase number (Venus fluorescence per cell) is plotted for: Tn4rev only strain: MG1655 Δlac pJK14-PLtet-01-Tn4rev & pZA31-PLlacOid-SmR (red, ◯), TnpB introduced in trans with Tn4rev: MG1655 Δlac pJK14-PLtet-01-Tn4rev pZA31-PLlacOid-mCherry-tnpB for different concentrations of IPTG (0μM (yellow, Δ), 10μM (green, ☐), 20μM (turquoise, ✳), 50μM (teal, ▽), 100μM (navy, ◁), 200μM (magenta, ✋) & 2000μM (violet, ✲)), and TnpB introduced in cis with Tn4rev: MG1655 Δlac pJK14-PLtet-01-Tn4rev-mCherry-tnpB & pZA31-PLlacOid-SmR (orange, X). Both Venus-TnpA concentration and the excision rate of TEs from plasmids increase with TnpB. As the concentration of transposase increases, the rate of excision events increases proportionally until it reaches its peak value.

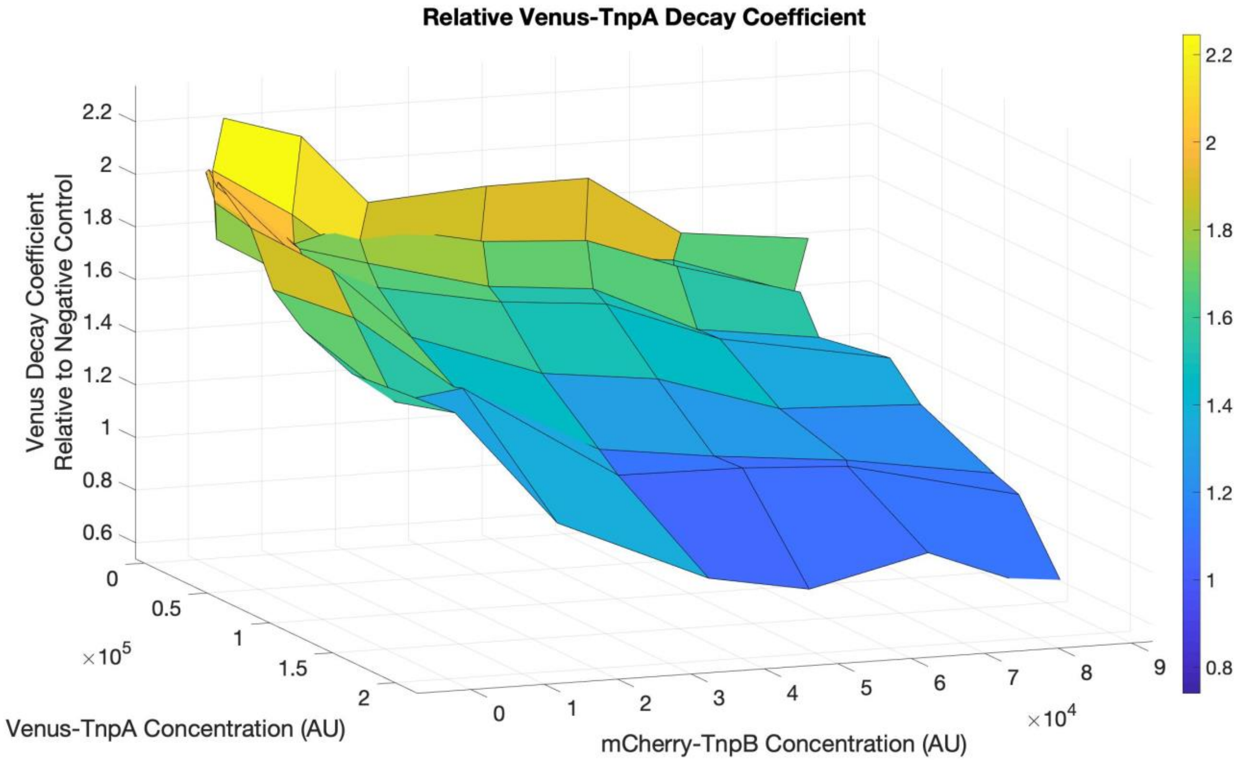


**Figure S6: Decay of Number of Transpose Molecules per cell over time:** The decay coefficients of the transposase concentration over time, $\beta g$, for the strain with introduction of TnpB *in trans* with Tn4rev are normalized to values for the negative control strain and plotted for all concentrations of TnpA and TnpB are plotted. For lower concentrations of both proteins, the number of transposase molecules decays at a higher rate over time. Increase in TnpA and TnpB both result in maintenance of transposase concentration due to a lower decay rate over time.

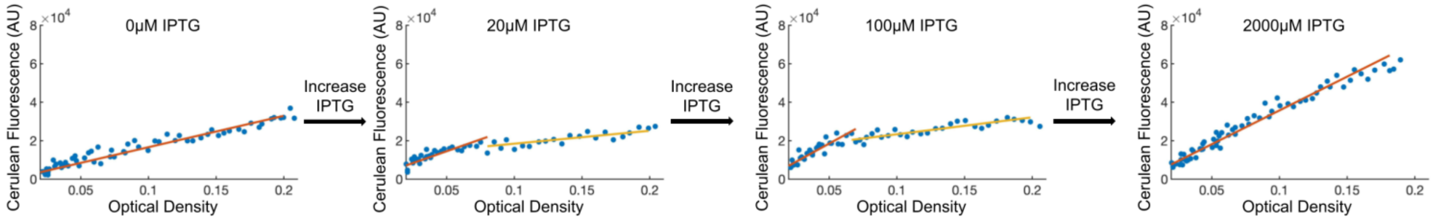


**Figure S7: Excision Rate Can Have Bimodal Slope During Exponential Growth.** Determination of average excision rate per cell as the slope of mCerulean3 fluorescence versus optical density. For intermediate IPTG concentrations (e.g., 20 μM and 100 μM shown here), the response can appear with bimodal slope. In such cases, the integral of fluorescence versus OD (Cumulative Cerulean) is calculated as given in eq. S3.2.3, with $m_1$ and $m_2$ determined as the slopes of the red and yellow lines, respectively.

#### SUPPLEMENTAL TABLES

**Table S1. Quantitative Features of Venus-TnpA Induction Response**

| **Strain** | **Peak Venus Fluorescence (AU)** | **Inflection Point (aTc)** | **Slope at Inflection Point (AU)** |
|---|---|---|---|
| **Tn4rev Only** | **1.15± 0.12 x $10^5$** | **64.29 ± 11.92** | **1.63 ± 0.25 x $10^5$** |
| **TnpB *in cis* Tn4rev** | **1.74± 0.10 x $10^5$** | **28.75 ± 15.93** | **2.27 ± 1.10 x $10^5$** |
| **TnpB i*n trans* Tn4rev (No IPTG)** | **1.19± 0.12 x $10^5$** | **64.25 ± 17.39** | **1.71 ± 0.37 x $10^5$** |
| **Immobilized TE -TnpB -IPTG** | **1.81 ± 0.17 x $10^5$** | **7.98 ± 1.90** | **3.14 ± 1.10 x $10^5$** |
| **Immobilized TE -TnpB +IPTG** | **1.68 ± 0.24 x $10^5$** | **9.60 ± 1.25** | **3.85 ± 1.34 x $10^5$** |
| **Immobilized TE +TnpB -IPTG** | **1.26 ± 0.11 x $10^5$** | **8.35 ± 0.74** | **2.22 ± 0.29 x $10^5$** |
| **Immobilized TE +TnpB +IPTG** | **1.31 ± 0.09 x $10^5$** | **6.86 ± 0.60** | **2.36 ± 0.34 x $10^5$** |

**Table S2. Quantitative Features of Excision Rate Response**

| **Strain** | **Peak Cerulean Fluorescence (AU)** | **Inflection Point (aTc)** | **Slope at Inflection Point (AU)** |
|---|---|---|---|
| **TnpB *in cis* Tn4rev** | **$2.01 \pm 0.10 \times 10^5$** | **$24.99 \pm 1.80$** | **$3.75 \pm 0.39 \times 10^5$** |

**Table S3: Quantitative Features of Excision Number Response**

| **Strain** | **Peak Cumulative Cerulean Fluorescence (AU)** | **Inflection Point (aTc)** | **Slope at Inflection Point (AU)** |
|---|---|---|---|
| **TnpB *in cis* Tn4rev** | **$1.17 \pm 0.10 \times 10^8$** | **$23.69 \pm 1.43$** | **$1.42 \pm 0.15 \times 10^8$** |

**Table S4: P-values for Changes in Total Transposon Number**

| Transition (growth with) | | P-value | Significance |
|---|---|---|---|
| No TnpA and no TnpB | With TnpA and no TnpB | < 0.00001 | Yes |
| With TnpA and no TnpB | With TnpA and with TnpB | 0.00022 | Yes |
| No TnpA and no TnpB | With TnpA and with TnpB | 0.0499 | No |

**Table S5: P-values for Changes in Plasmid-Transposon Number**

| Transition (growth with) | | P-value | Significance |
|---|---|---|---|
| No TnpA and no TnpB | With TnpA and no TnpB | < 0.00001 | Yes |
| With TnpA and no TnpB | With TnpA and with TnpB | 0.004 | Yes |
| No TnpA and no TnpB | With TnpA and with TnpB | < 0.00001 | Yes |

**Table S6: Quality of Fit of Exponential Decay of Growth Rate**

| Strain | Tn4rev Only | *In trans*, 0μM IPTG | 10μM IPTG | 20μM IPTG | 50μM IPTG | 100μM IPTG | 200μM IPTG | 2mM IPTG | *In cis* |
|---|---|---|---|---|---|---|---|---|---|
| $R^2$ value | 0.9244 | 0.411 | 0.9347 | 0.9527 | 0.9385 | 0.9243 | 0.8122 | 0.9601 | 0.9498 |

**Table S7: Primer Sequences for qPCR**

| Reaction Number | Amplicon | Primer Pairs | Sequence |
|---|---|---|---|
| 1 | *nth* gene of MG1655 for cell number | MG1655-nth F | 5'-GATCCTCACTCGCCTGCGTGAGAAC-3' |
| | | MG1655-nth R | 5'-CAAGCATCGCTGCAGGCGTATTCG-3' |
| 2 | *mCerulean* gene for plasmid copy number | DK-pJK14-qPCR F | 5'-CAAGTAATCACCGTTTGGACCTTGGG-3' |
| | | DK-pJK14-qPCR R | 5'-GAAAGGGCAGATTGTGTGGACAGG-3' |
| 3 | *venus* gene for transposon number | *tnpA-venus*-qPCR F | 5'-GTTACTCATTCCTCCTCCTCCTCCC-3' |
| | | *tnpA-venus*-qPCR R | 5'-CAACCACTACCTGAGCTACCAGTCC-3' |
| 4 | Plasmid-Tn4rev joint for excision rate | pJK14-RE\IP-qPCR F | 5'-CGCATTTACGTTGACACCACCTTTGAC-3' |
| | | pJK14-RE\IP-qPCR R | 5'-CTCGCTGATAATTGTGAGCGCTCAC-3' |

(1) Kim Neil H.; Lee Gloria; Sherer Nicholas A.; Martini K. Michael; Goldenfeld Nigel; Kuhlman Thomas E. Real-Time Transposable Element Activity in Individual Live Cells. *Proc. Natl. Acad. Sci.* **2016**, *113* (26), 7278–7283. https://doi.org/10.1073/pnas.1601833113.

(2) Gene Location and DNA Density Determine Transcription Factor Distributions in Escherichia Coli. *Mol. Syst. Biol.* **2012**, *8* (1), 610. https://doi.org/10.1038/msb.2012.42.

(3) Kuhlman, T. E.; Cox, E. C. Site-Specific Chromosomal Integration of Large Synthetic Constructs. *Nucleic Acids Res.* **2010**, *38* (6), e92. https://doi.org/10.1093/nar/gkp1193.

(4) Tas, H.; Nguyen, C. T.; Patel, R.; Kim, N. H.; Kuhlman, T. E. An Integrated System for Precise Genome Modification in Escherichia Coli. *PloS One* **2015**, *10* (9), e0136963. https://doi.org/10.1371/journal.pone.0136963.
